# Chromosome-arm-specific telomere length governs dual modes of structural genome evolution in IDH-mutant astrocytoma

**DOI:** 10.64898/2026.04.22.720226

**Authors:** Maryam Jehangir, Kristen L. Drucker, Ayse Keskus, T. Rhyker Ranallo-Benavidez, Orlena Benamozig, Thomas Kollmeyer, Terry C. Burns, Caterina Giannini, Benjamin R. Kipp, Stephen J. Murphy, Amber R. Bridgeman, Jesse R. Walsh, Ramanath Majumdar, Daniel Lachance, Jeanette Eckel-Passow, Ofer Shoshani, Mikhail Kolmogorov, Robert B. Jenkins, Floris P. Barthel

## Abstract

IDH-mutant astrocytomas maintain telomeres through the alternative lengthening of telomeres (ALT) pathway, producing extreme inter-arm telomere length heterogeneity, yet how this heterogeneity shapes structural genome evolution remains unknown. Using Oxford Nanopore long-read sequencing of 20 IDH-mutant astrocytomas, we profiled structural variants (SVs), copy number variants, extrachromosomal DNA (ecDNA) and measured allele-specific telomere lengths from individual long reads. We identified pervasive complex rearrangements, including chromothripsis and foldback events consistent with breakage-fusion-bridge cycles, and widespread ecDNAs. SV breakpoints were enriched at telomeric and centromeric regions regardless of local telomere length, revealing constitutive structural fragility. Arm-level telomere length analysis uncovered a dual-mode model: arms with short telomeres preferentially harbored breakage-associated events, while arms with long ALT-maintained telomeres were enriched for ecDNA and amplification-associated events. These findings identify chromosome-arm-specific telomere length as a determinant of structural genome evolution in ALT-driven tumors.

## INTRODUCTION

Diffuse gliomas comprise a heterogeneous group of primary brain tumors whose genomic landscape is defined by discrete molecular subtypes with distinct diagnostic, prognostic, and therapeutic implications (Barthel, Johnson, Wesseling, & Verhaak, 2018; Cancer Genome Atlas Research et al., 2015; Ceccarelli et al., 2016; Eckel-Passow et al., 2015). Among these, isocitrate dehydrogenase (IDH)-mutant astrocytomas represent a biologically distinct tumor entity, distinguished by a unique pattern of structural variation (SV) and copy number variation (CNV). Unlike IDH wild-type glioblastoma, which harbors frequent arm-level aneuploidy events (e.g., gain of chromosome 7 and loss of chromosome 10), or IDH-mutant oligodendroglioma, which is defined by the combined loss of chromosome arms 1p and 19q, astrocytomas accumulate abundant segmental CNVs and complex SVs rather than whole-arm or whole-chromosome alterations (Cancer Genome Atlas Research et al., 2015; Ceccarelli et al., 2016; Drucker et al., 2026). Longitudinal studies have demonstrated that this genomic instability is an active and ongoing process: recurrent IDH-mutant astrocytomas exhibit a substantial increase in CNV burden compared to their matched primary tumors, contrasting with glioblastoma, where foundational genotypes appear largely established prior to clinical presentation (Barthel et al., 2019; Barthel, Wesseling, & Verhaak, 2018; Varn et al., 2022).

Telomere maintenance is intimately linked to the biology of diffuse gliomas and may fundamentally shape the trajectory of genomic instability. Glioblastomas and oligodendrogliomas harbor activating mutations in the telomerase reverse transcriptase *TERT* promoter, constitutively reactivating telomerase to counteract telomere attrition and stably maintain telomeres at a minimal functional length (Eckel-Passow et al., 2015; Killela et al., 2013). IDH-mutant astrocytomas follow a fundamentally different strategy. These tumors carry mutations in *TP53* and *ATRX*, lack *TERT* expression, and instead maintain their telomeres through the alternative lengthening of telomeres (ALT) pathway, a recombination-based mechanism driven by break-induced replication (BIR) at telomeric ends (Heaphy, de Wilde, et al., 2011; Lovejoy et al., 2012; Roumelioti et al., 2016; Zhang, Yadav, Ouyang, Lan, & Zou, 2019). In telomeric BIR, unprotected telomere ends invade homologous templates, such as a sister chromatid telomere or extrachromosomal telomeric DNA, to initiate long-range DNA synthesis and extend the telomere (Dilley et al., 2016). This process yields telomeres that are remarkably long and heterogeneous in length, a hallmark feature of ALT-positive tumors across cancer types (Barthel et al., 2017; Heaphy, Subhawong, et al., 2011). The inherent instability of ALT-mediated telomere maintenance, characterized by cycles of erosion, recombination, and heterogeneous elongation, raises the possibility that chromosome-specific telomere length directly influences the structural fate of individual chromosome arms.

A parallel feature of glioma genomes is the amplification of oncogenes on extrachromosomal DNA (ecDNA), circular elements linked to intratumoral heterogeneity, aggressive disease, and treatment resistance across cancer types (Kim et al., 2020; Shoshani et al., 2021; Yi, Chamorro Gonzalez, Henssen, & Verhaak, 2022). ecDNA biogenesis is mechanistically coupled to chromothripsis and breakage-fusion-bridge (BFB) cycles, both of which can be initiated by telomere dysfunction. Critically short or uncapped telomeres generate dicentric chromosomes whose breakage through successive cell divisions initiates BFB cycles, chromothripsis, and the formation of micronuclei harboring fragmented chromosomes (Maciejowski et al., 2020; Maciejowski, Li, Bosco, Campbell, & de Lange, 2015; Umbreit et al., 2020). Chromosome mis-segregation into micronuclei, whether from bridge breakage or other mitotic errors, triggers additional rounds of shattering that can generate ecDNA de novo (Ly et al., 2019; Shoshani et al., 2021). While ecDNA is a well-characterized feature of IDH wild-type glioblastoma, its prevalence and genomic context in IDH-mutant astrocytoma remain poorly understood, and whether chromosome-arm-specific telomere length governs ecDNA formation in ALT-driven tumors has not been investigated (deCarvalho et al., 2018; Noorani et al., 2025). Moreover, although large-scale genomic studies of diffuse glioma have catalogued arm-level aneuploidy and copy number patterns across subtypes and time (Barthel et al., 2019; Ceccarelli et al., 2016; Drucker et al., 2026), the relationship between these structural features and telomere biology has not been systematically examined.

Here, we sought to define the genomic correlates of chromosome-arm-level telomere length, including SVs, CNVs, and ecDNA, in ALT-driven IDH-mutant astrocytomas. We first contextualized the structural genomic landscape of IDH-mutant astrocytomas relative to other glioma subtypes using TCGA and GLASS cohort data (n=760). We then performed Oxford Nanopore Technologies (ONT) long-read sequencing on 20 fresh-frozen primary IDH-mutant astrocytoma tumor samples (Figure 1A), comprehensively profiling these features of genomic instability at chromosome-arm resolution. We specifically asked whether SVs are enriched near telomeric and centromeric regions, whether chromosome-arm-specific telomere length variability is associated with local SV and CNV burden, and whether ecDNA formation is linked to telomere state. Our analyses revealed a heterogeneous landscape of complex somatic SVs, with breakpoints significantly enriched at telomeric and centromeric regions. We identified widespread ecDNA across the cohort, with both chromothripsis and ecDNA burden associated with advanced tumor grade. Arm-level telomere length analysis revealed that breakage-associated variants, including chromothripsis, were enriched on arms with shorter telomeres, while amplification-associated features, including ecDNA, were enriched on arms with longer telomeres. These findings establish the structural complexity of ALT-driven astrocytoma genomes and identify telomere-proximal regions as hotspots for structural rearrangement.

**Figure 1.**
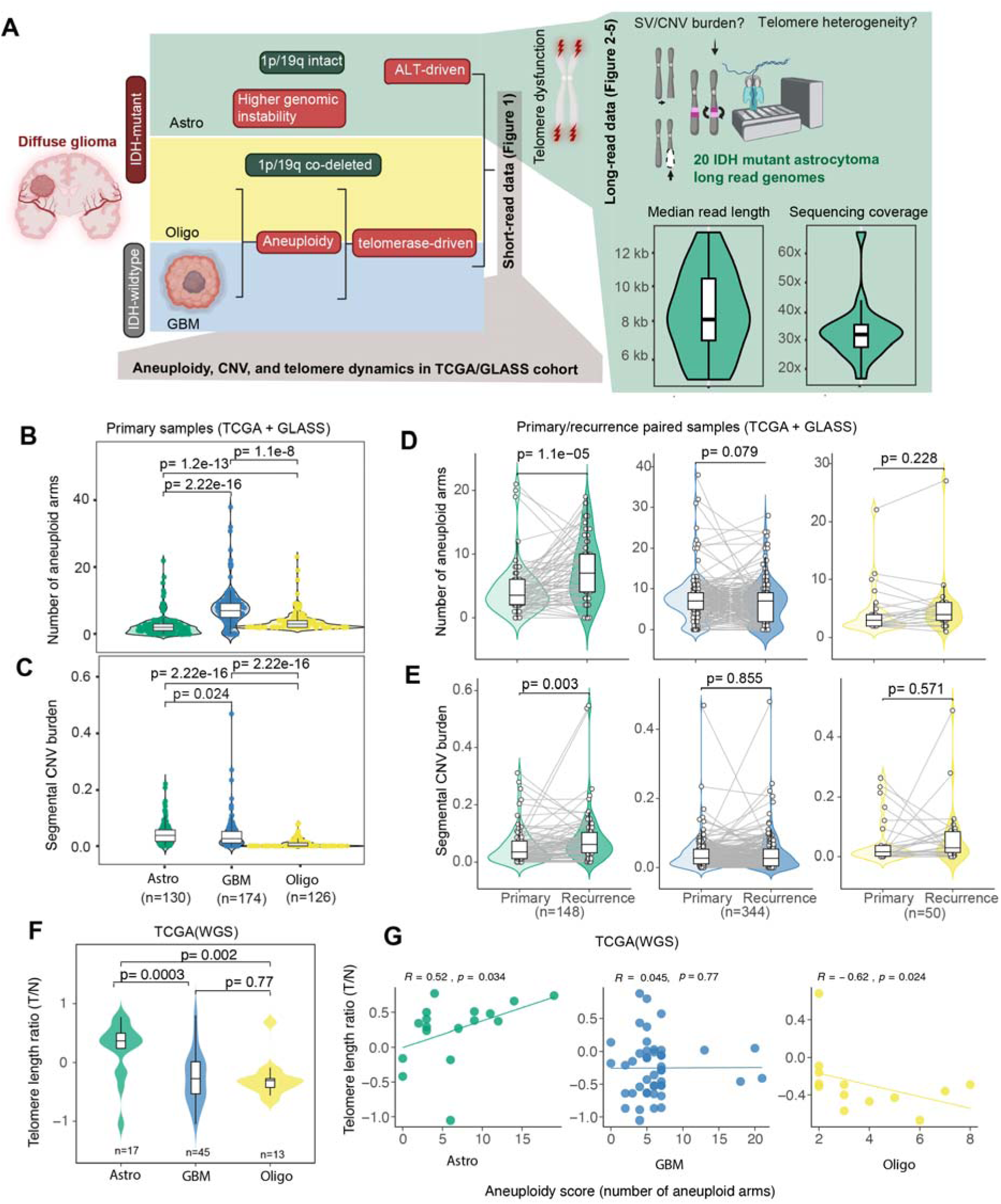
Astrocytoma genomes featured high level genomic instability and telomere heterogeneity. **(A)** A schematic view of genomic characteristics in diffuse gliomas, and workflow of investigating the complex mechanisms of astrocytoma genomes using long-read sequencing created using BioRender. **(B–E)** Quantitative comparison of aneuploidy and segmental copy number variation (CNV) burden across molecular glioma subtypes Astrocytomas (Astro)(green), glioblastomas (GBM) (blue), and oligodendrogliomas (Oligo)(yellow) are shown, with data separated into primary and recurrent tumor groups. Each point represents an individual tumor sample, and boxplots indicate median and interquartile range. Statistical significance was assessed using the Wilcoxon rank-sum test for unpaired samples and the Wilcoxon signed-rank test for paired comparisons. **(F)** Distribution of tumor-to-matched-normal (T/N) telomere content ratios across diffuse glioma subtypes, revealing significantly higher T/N telomere content ratios in astrocytomas compared with glioblastomas and oligodendrogliomas (Wilcoxon rank-sum test). **(G)** Spearman correlation analyses between telomere T/N ratio and aneuploidy burden highlight subtype-specific relationships between telomere state and aneuploidy in astrocytomas, glioblastomas and oligodendrogliomas.

## RESULTS

### Astrocytoma genomes are defined by segmental aneuploidy and variable telomere length dynamics

We surveyed the TCGA and GLASS cohorts (n = 760) to characterize the structural genomic differences comparing IDH-mutant astrocytomas to other glioma subtypes (Figure 1A). Glioblastomas and oligodendrogliomas exhibited a high burden of arm-level aneuploidy compared to astrocytomas (Figure 1B). Despite the lower frequency of arm-level aneuploidy, IDH-mutant astrocytomas carried a significantly higher burden of segmental CNVs than glioblastoma or oligodendroglioma, consistent with a mechanistically distinct origin of copy number change (Figure 1C). Longitudinal comparison revealed that astrocytomas undergo a significant increase in genomic instability from primary to recurrence, both in terms of aneuploidy and segmental CNVs, implying that unlike glioblastoma, astrocytoma genomes are structurally evolving over the course of disease (Figure 1D–E). These alterations clustered on discrete chromosome arms (Figure S1A), indicating that structural remodeling preferentially targets specific arms rather than occurring uniformly across all chromosomes, consistent with prior observations linking arm-level CNVs to specific oncogenes and tumor suppressor genes (Drucker et al., 2026; Shih et al., 2023). Arm-level aneuploidy analysis comparing primary to recurrent astrocytomas revealed that 10q, 4p, 4q, 7p and 7q were more likely to be affected by *de novo* aneuploidy events than other chromosome arms (Figure S1A).

Astrocytomas were further distinguished by unique telomere dynamics, exhibiting a significantly higher tumor-to-normal telomere content ratio than glioblastoma and oligodendroglioma (Figure 1F). Telomere content in astrocytoma was higher on average and more dispersed than in glioblastoma or oligodendroglioma, with heterogeneous telomere elongation and shortening consistent with telomere maintenance by the ALT pathway (Figure 1F, Figure S1B), in alignment with prior data (Barthel et al., 2017).

This heightened telomeric variability coincided with an increased segmental CNV burden and large-scale genomic instability. Notably, in astrocytomas, higher tumor-to-normal telomere content ratios correlated positively with both the number of aneuploid arms and the segmental CNV burden, consistent with a role for telomere length in genome remodeling (Figure 1G; Figure S1C). Taken together, these data indicate that telomere length and segmental aneuploidy are tightly associated variables in IDH mutant astrocytoma.

### Long-read sequencing identifies abundant complex structural variants in a cohort of astrocytomas

To resolve the interplay between structural variants (SVs), copy number variants (CNVs), and telomere length at chromosome-arm resolution, we performed long-read Oxford Nanopore (ONT) sequencing on tumors from n = 20 patients with molecular testing and pathology review confirmed IDH-mutant astrocytoma (n = 7 grade 2, n = 13 grade 3; Figure 1A). Clinical copy-number profiling using Oncoscan and Illumina EPIC arrays and mutation analysis with directed sequencing revealed a high frequency of recurrent genomic alterations, with TP53 (65%; 13/20) and ATRX (76.47%; 13/17) being the most frequently altered genes. CDKN2A/B was intact in at least one copy across all samples, consistent with none of the tumors reaching grade 4 (Figure 2A, Supplementary Table 1). Arm-level aneuploidy was detected in 65% (n = 14/20) of tumors (Supplementary Table 2). Long-read data yielded median read lengths exceeding 12 kb, and ∼30× genome-wide coverage (Figure 2B, Supplementary Figure S2A; Supplementary Table 3), with alignment rates to the T2T-CHM13 v2.0 reference exceeding 95% in all cases (Nurk et al., 2022).

**Figure 2.**
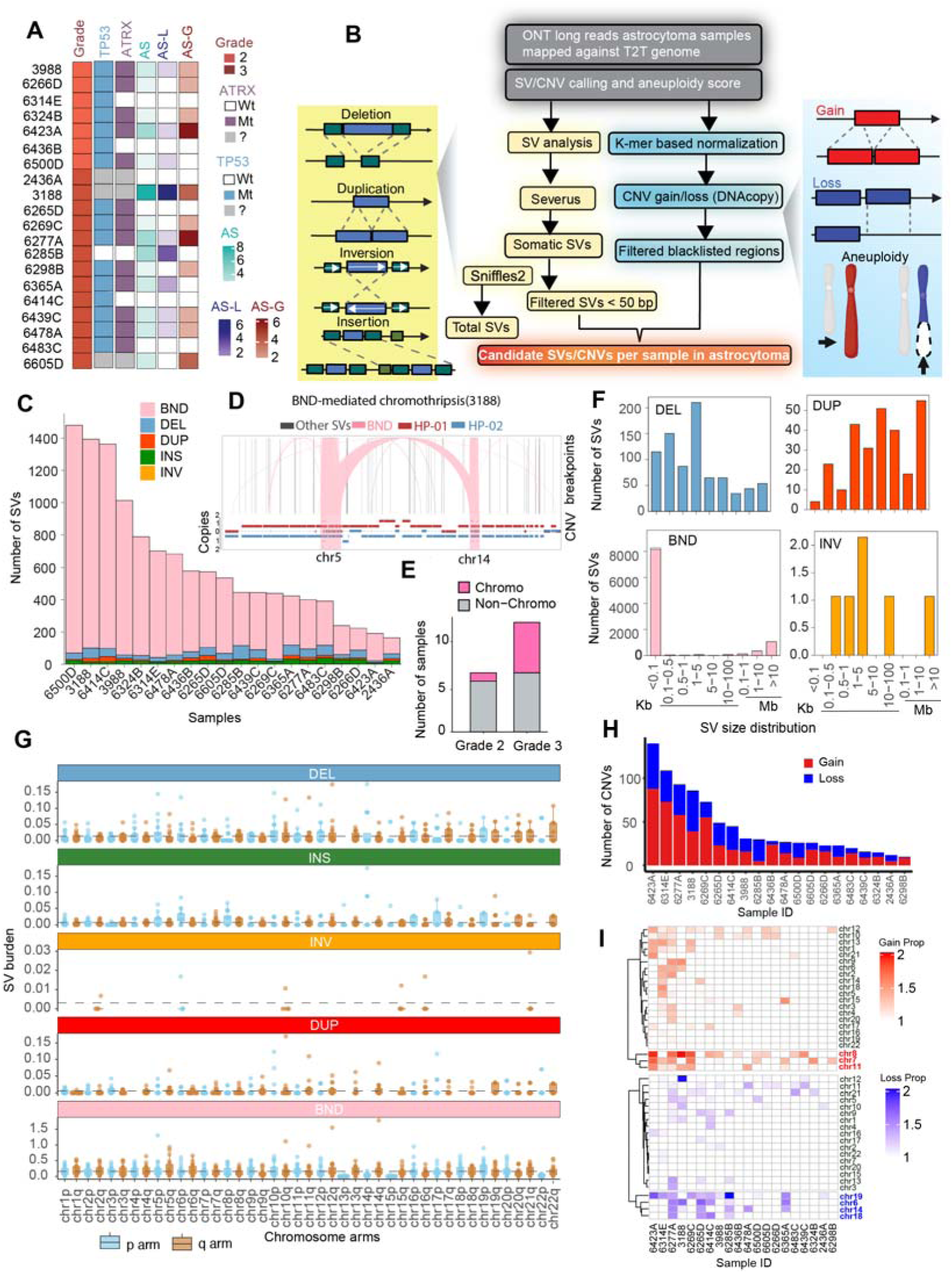
Long-read sequencing defines the structural variant landscape of IDH-mutant astrocytomas. **(A)** Heatmap summarizing clinical, molecular, and genomic features across 20 astrocytoma samples sequenced using Oxford Nanopore Technologies (ONT). Samples are sorted by WHO grade (grade 2 followed by grade 3). Shown are key clinical annotations, oncogenic drivers (e.g., *TP53* and *ATRX* mutations), and chromosome-arm–level aneuploidy profiles. Chromosome-arm-level aneuploidy was quantified using the Aneuploidy Score (AS), with separate metrics for gains (AS-G) and losses (AS-L), representing the number of chromosome arms harboring copy-number gains or losses, respectively. **(B)** Workflow schematic outlining the variant detection pipeline, including alignment to the CHM13 reference genome, germline-inclusive SV calling using Sniffles2, somatic SV detection using Severus and CNV profiling. **(C)** Stacked bar plots showing per-sample counts of SV types, classified as insertions (green), deletions (blue), duplications (red), inversions (orange), and translocations/BNDs (pink). **(D**) Representative chromothripsis event from sample (3188) in the manually edited Wakhan output. Pink BND links connect two chromosomes (chr5 and 14) and form a dense cluster of breakpoints within a restricted region. The phased copy-number tracks (hap1, red; hap2, blue) show matching oscillating copy-number states across the same interval, supporting chromothripsis by both breakpoint clustering and the CNV pattern. **(E)** Barplot showing the number of samples with and without chromothripsis. **(F)** SV size distributions across variant classes, demonstrating that deletions typically span kilobase to low-megabase ranges, whereas duplications, inversions, and translocations represent larger rearrangements. Colors correspond to SV types as in (C). **(G)** Cumulative SV burden per chromosome arm across the cohort, with each point representing a chromosome arm per sample. The y-axis denotes SV density (Mbl□^1^), and p- and q-arms are colored blue and brown, respectively**. (H)** Sample-wise CNV profiles showing that copy-number gains (red) are more frequent but smaller in size, whereas losses (blue) are less frequent but span larger genomic regions. **(I)** Chromosome-level heatmaps depicting CNV burden across samples, where color intensity represents the fraction of each chromosome affected, providing an integrated view of genome-wide copy-number variation in astrocytomas. In both **(H)** and **(I)**, samples are sorted by total CNV counts, while chromosomes (rows) are hierarchically clustered based on CNV profile similarity to highlight recurrent patterns of genomic instability.

We identified somatic SVs using Severus (Keskus et al., 2025), which subtracts germline variants using population panels and resolves complex rearrangement structures from breakend (BND) clusters, and called germline-inclusive SVs with Sniffles2 (Smolka et al., 2024). We detected a mean of 550 somatic SVs per tumor (range 164–1,479; Figure 2C, Figure S2B–C, Supplementary Table 4), revealing extensive inter-tumor heterogeneity. Complex inversions comprised a large fraction of events in high-burden tumors (approaching ∼60%), frequently accompanied by tandem duplications (Figure S2C). Integration with Wakhan for haplotype-resolved CNV phasing enabled classification of diverse rearrangement signatures, including chromothripsis, chromoplexy, and foldback inversion-associated translocations (Figure S3; Supplementary Table 5) (Ahmad et al., 2025). For example, tumor 3188 harbored a multi-fragment inter-chromosomal rearrangement linking chromosomes 5 and 14, consistent with chromothripsis (Figure 2D), while tumor 6277A exhibited a chromoplexy-like configuration involving five chromosomes (Supplementary Table 5). Chromothripsis was detected in 7 of 20 tumors (35%), affecting 17 chromosome arms across the cohort. Chromothripsis-bearing tumors were predominantly grade 3 (6/7 86%), compared with 1/7 (14%) grade 2 tumors, though this enrichment did not reach statistical significance (Fisher’s exact test, OR = 5.14, p = 0.33; Figure 2E), likely reflecting the limited sample size. Somatic SVs ranged from a few kilobases to over 10 Mb, with inversions and duplications spanning the largest genomic intervals (Figure 2F).

At chromosome-arm resolution, somatic SV burden revealed distinctive distribution patterns (Figure 2G). To quantify how unevenly SVs are distributed across arms, we applied the Gini index, a measure of dispersion (0 = perfectly uniform distribution across arms; 1 = all events concentrated on a single arm), alongside prevalence, defined as the fraction of sample-arm observations with non-zero SV burden. BND-type events were the most broadly distributed class (Gini = 0.53, prevalence = 88%), while deletions showed intermediate concentration (Gini = 0.72, prevalence = 47%). Insertions and duplications were markedly more concentrated (Gini = 0.78 and 0.89, respectively), with burden confined to a minority of chromosome arms (prevalence = 34% and 19%; Figure 2G, Figure S2D). Reciprocal inversions were too rare for meaningful Gini estimation (6 events across the cohort). These patterns indicate that while BND-type events are dispersed features of astrocytoma genomes, deletions, insertions, and duplications cluster in progressively smaller subsets of hotspot arms. By contrast, Gini analysis of the Sniffles2 data, representing the combination of germline and somatic variants (mean, 29.9k SVs per sample; Fig. S4A) showed a near-uniform distribution for deletions and insertions (Gini = 0.27 and 0.24, prevalence = 98% each), with the modest departure from uniformity driven primarily by enrichment on acrocentric p-arms (chr14p, chr22p, chr21p; 1.7–2.2x higher burden), consistent with known polymorphism in ribosomal DNA and satellite-rich arrays (Fig. S4B–E). Germline-inclusive duplications and inversions were more concentrated (Gini = 0.77 and 0.81), consistent with known segmental duplication and inversion polymorphism hotspots (Fig. S4D).

CNV profiling using DNAcopy revealed that tumors carried an average of 50 segmental CNVs, with gains more frequent but smaller (median ∼25 Mb) and losses less frequent but larger (median ∼100 Mb; Figure 2H, Supplementary Figure S5A–B) (Olshen, Venkatraman, Lucito, & Wigler, 2004). Recurrent segmental losses affected chromosomes 5, 14, and 19, while gains recurred on chromosomes 7 and 8. Per-chromosome and per-sample analysis (Figure 2I; Supplementary Figure S5C) revealed substantial heterogeneity: some tumors accumulated abundant segmental events, while others were dominated by arm-level alterations, suggesting at least two distinct modes of genome remodeling. Together, these data establish the structurally complex genomic landscape of IDH-mutant astrocytomas and provide a foundation for interrogating how telomere dynamics shape ongoing genome instability.

### Extrachromosomal DNA is pervasive in IDH-mutant astrocytomas and associates with tumor grade

Extrachromosomal DNA (ecDNA) is a well-characterized feature of IDH wild-type glioblastoma, but its prevalence in IDH-mutant astrocytoma has not been systematically assessed (deCarvalho et al., 2018; Kim et al., 2020; Noorani et al., 2025). To separately characterize ecDNA and focal gain events in our cohort, we performed circular DNA assembly using Decoil to detect ecDNA and identified highly gained regions (HGs) from CNV segments exceeding a stringent copy-number threshold (Figure 3A; see Methods) (Giurgiu et al., 2024).

**Figure 3.**
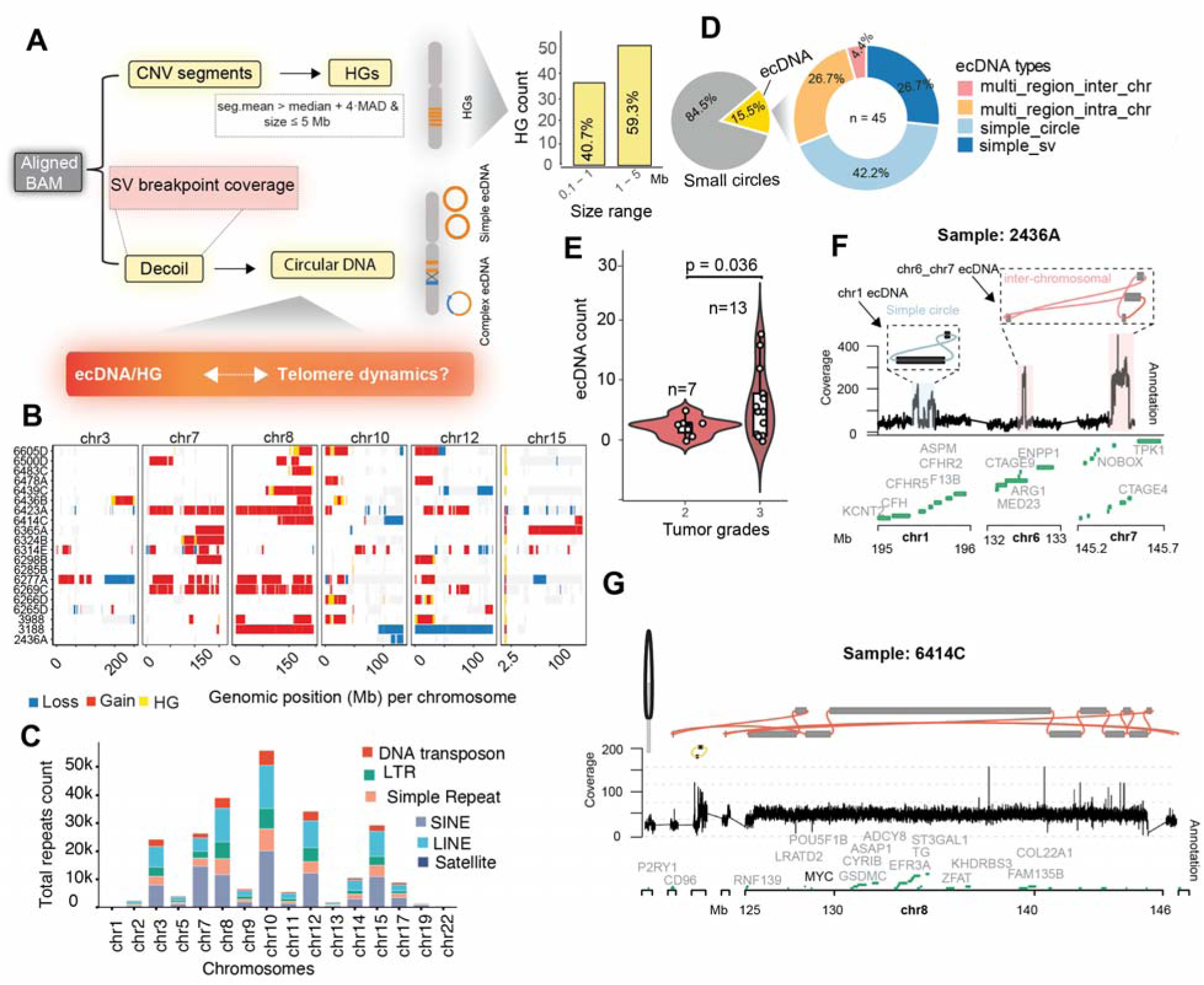
Extrachromosomal DNA is pervasive in IDH-mutant astrocytomas and associates with tumor grade. **(A)** Workflow schematic illustrating detection of ecDNA and highly gained regions (HGs) using the Decoil pipeline on long-read sequencing data. **(B)** Heatmap displaying CNV segment values restricted to chromosomes harboring a high burden of HGs. Rows represent genomic coordinates across selected chromosomes and columns represent individual samples. HGs appear as sharp yellow focal peaks. Moderate gains (log_2_ > +0.2) and losses (log_2_< –0.2) are shown in red and blue, respectively. Recurrent HGs emerge at conserved chromosomal coordinates, indicating hotspot loci subjected to intense focal amplification pressure **(C)** Comprehensive distribution of major repeat element families (SINE, LINE, LTR, DNA transposon, and Satellite) overlapping HGs. Chromosomes are ordered numerically to illustrate the global repetitive landscape associated with focal amplifications. **(D)** Pie charts summarizing circular DNA calls across the cohort. Left: distribution of all 290 circular DNA elements into canonical ecDNA (n = 45) and short circular DNAs (n = 245). Right: topology distribution of canonical ecDNA elements, with consistent colors as in subsequent panels. Percentages are indicated. **(E)** Association of ecDNA counts with astrocytoma grade; each dot represents a sample, showing significantly higher ecDNA burden in grade 3 (dark red) compared with grade 2 (light red) tumors. Statistical significance was determined using a t-test. **(F-G)** Representative ecDNA configurations were reconstructed using Decoil, illustrating the structural heterogeneity observed across the astrocytoma cohort. These examples encompass a spectrum of complexity, ranging from simple intrachromosomal circularizations to highly rearranged multi-segment or inter-chromosomal configurations**. (F)** In sample 2436A, a chimeric multi-chromosomal assembly is depicted, involving segments from chromosomes 6 and 7 that harbor oncogenic drivers. **(G)** At the 8q24.21 locus, a complex intrachromosomal ecDNA highlights massive focal amplification of the *MYC* oncogene. In both panels, coverage tracks (top) reveal high-magnitude focal amplification across the amplicons, while gene annotations (middle) delineate coding sequences encompassed within each structure. Connecting arcs (bottom) depict inferred breakpoint junctions and circular topology, with colors of ecDNA subtypes match those used in the pie chart.

We identified 61 HGs (Methods, briefly defined as < 5 Mb and > 4 median absolute deviations over the sample median copy number) across 16 chromosomes in 13 of 20 samples, with sizes ranging from 0.1 to 5 Mb (41% below 1 Mb, 59% between 1–5 Mb; Figure 3A–B, Figure S6A). HGs were non-randomly distributed, with recurrent enrichment on chromosomes 7, 8, 1, and 12 (Figure 3B; Figure S6B). Recurrently amplified loci harbored oncogenes such as *EZH2*, *KDM5A*, *ZEB1*, and *SOX9* (Figure S6C, Supplementary Table 6) and overlapped a repeat-rich landscape dominated by retroelements (Figure 3C).

To determine whether focally gained regions could represent circular elements, we reconstructed circular DNA structures using Decoil, identifying *n* = 290 unique circular DNA elements across 19 tumors (Methods). These could be further broken down into 45 canonical ecDNA elements (>100 kb) and 245 shorter circular DNAs (Figure 3D; Supplementary Table 7). Among the 45 canonical ecDNA elements, topologies were more diverse: simple circles (42%), simple structural variant-associated (27%), multi-region intra-chromosomal (27%), and multi-region inter-chromosomal (4%). Consistent with this architectural diversity, ecDNA segments were distributed unevenly across the genome, with enrichment on chromosomes 7, 1, 8, and 17, and showed marked intertumoral variability in burden (Supplementary Figure 7A–C). Canonical ecDNA elements harbored coding sequences of known oncogenes including *MYC*, *CCN4*, and *ASPM* (Supplementary Table 6; Figure S7D). Annotation of ecDNA intervals further showed that these structures contain a range of tandem (satellites and simple repeats) and interspersed (transposable elements) repetitive sequence, predominantly retroelements including long interspersed nuclear elements (LINEs), short interspersed nuclear elements (SINEs), long terminal repeat (LTR) elements and DNA transposons. Among these, SINEs and LINEs were the most abundant, representing 32% and 28% of the total repetitive sequence fraction (Figure S7E). Grade 3 astrocytomas carried significantly more ecDNA than grade 2 tumors, indicating that ecDNA accumulation accompanies tumor progression (Figure 3E). ecDNA assemblies exhibited structurally diverse configurations, ranging from simple single-chromosome circles to complex multi-chromosome clusters with variable gene content (Figure 3F-G; Figure S7F).

### Chromosome-arm-specific telomere length analysis reveals heterogeneity consistent with ALT and dual modes of genomic instability

Given the structural diversity of IDH-mutant astrocytomas and the inherent heterogeneity of ALT-mediated telomere maintenance, we asked whether chromosome-arm-specific telomere length (TL) is associated with the type and distribution of SVs and CNVs across the genome. We estimated allele-specific telomere lengths using Telogator2 (Stephens, Ferrer, Boardman, Iyer, & Kocher, 2022; Stephens & Kocher, 2024), which identifies telomere-containing reads in long-read sequencing data, clusters them into distinct alleles based on telomere variant repeat (TVR) composition, and anchors each allele to a specific chromosome arm via subtelomeric alignment. For each allelic cluster, Telogator2 estimates telomere length as the 75th percentile of read lengths within that cluster. In a bulk tumor sample, multiple allelic clusters per arm may reflect distinct subclonal populations with different telomere lengths. We therefore derived three arm-level summary statistics across all allelic clusters: TL min (shortest cluster), TL max (longest cluster), and TL median (median cluster). These metrics enables us to ask whether different types of SVs and CNVs associate with specific allelic extremes (e.g., the shortest or longest telomere on a given arm) rather than only the arm-level average.

Arm-level TL median values spanned a broad range (0.1–13 kb), with substantial intra- and inter-tumor heterogeneity consistent with ALT-like telomere length distributions (Figure S8A-D; Supplementary Table 8). Globally z-scored telomere lengths (cohort mean = 4.97 kb, SD = 1.80 kb) revealed that while most chromosome arms retained near-normal telomere lengths (40.0%, z-score < 0.5), a subset exhibited critically short telomeres (2.7%, z-score < −1.5) and a long right tail of very long telomeres (7.6%, z-score > 1.5) characteristic of ALT (Figure 4A).

**Figure 4.**
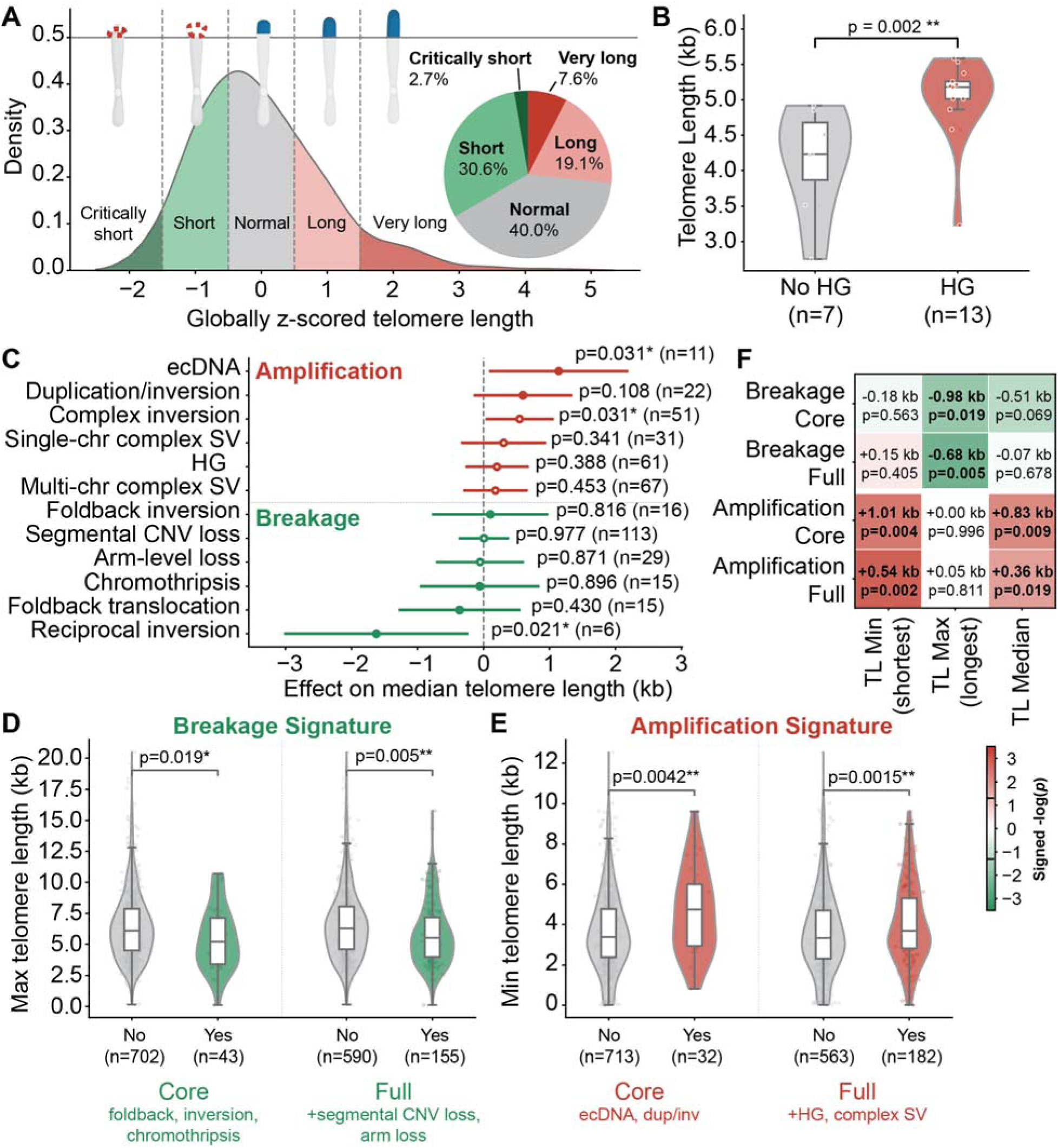
Telomere length drives dual modes of genomic instability in IDH-mutant astrocytoma. **(A)** Distribution of globally z-scored median telomere lengths across all chromosome arms in the cohort. Density plot (left) shows the z-score distribution in each telomere length category: critically short (z < -1.5; dark green), short (-1.5 to -0.5; light green), normal (-0.5 to 0.5; grey), long (0.5 to 1.5; light red), and very long (z > 1.5; dark red). Pie chart (right) shows the proportion of arms in each category. **(B)** Tumors harboring at least one HG or ecDNA event (n=13) exhibit significantly longer per-sample median telomere length than tumors without such events (n=7; Wilcoxon rank-sum p=0.002). **(C)** Forest plot of univariate mixed-effects model estimates for individual genomic events on median telomere length. Each point represents the within-patient effect (beta, in kb) of event presence versus absence, with 95% confidence intervals. Amplification-associated events (red) show positive effects (longer TL), while breakage-associated events (green) show negative effects (shorter TL). Filled circles indicate core signature components; open circles indicate secondary components. **(D)** Violin plots comparing maximum allelic telomere length (TL Max) between arms with and without the core (foldback inversions, reciprocal inversions, chromothripsis) and full (core + segmental CNV loss + arm-level aneuploidy loss) breakage signatures. P-values from linear mixed-effects models with patient as random intercept. **(E)** Violin plots comparing minimum allelic telomere length (TL Min) between arms with and without the core (multi-region intra-chromosomal ecDNA, duplicated inverted segments) and full (core + HG/ecDNA + complex inversions + complex rearrangements) amplification signatures. **(F)** Heatmap of mixed-effects model results testing all combinations of signature type (breakage core/full, amplification core/full) against three telomere length measures (TL Min, TL Max, TL Median). Cell color represents the signed -log10(p-value), with green indicating shorter TL on affected arms and red indicating longer TL. Effect sizes (beta, in kb) and p-values are annotated in each cell.

At the sample level, tumors harboring at least one HG event (n = 13) had significantly longer median telomere length than tumors without such events (n = 7; median TL 5.18 vs 4.23 kb; p = 0.002; Figure 4B), consistent with the possibility that ALT-driven telomere elongation and HG formation may be coupled. Visualization of arm-level telomere lengths overlaid with genomic event annotations further suggested that breakage-associated events (e.g., foldback inversions, chromothripsis) were concentrated on arms with shorter telomeres, while amplification-associated events (e.g., ecDNA, duplicated inversions) were concentrated on arms with longer telomeres (Supplementary Figure 9A–B), motivating formal statistical testing.

Univariate mixed-effects models testing individual genomic events against arm-level TL median revealed a divergence in the direction of association (Figure 4C; Supplementary Table 9). Amplification-associated events showed positive effects on telomere length: multi-region intra-chromosomal ecDNA had the strongest association (β = +1.14 kb, p = 0.031), followed by complex inversions (β = +0.55 kb, p = 0.031). Breakage-associated events generally showed weaker and non-significant effects on TL Median at the individual event level: foldback translocations (β = −0.36 kb, p = 0.430), chromothripsis (β = −0.06 kb, p = 0.896), and reciprocal inversions (β = −1.62 kb, p = 0.021; n = 6 arms) trended toward or reached shorter telomeres, though most individual breakage event types were too sparse to reach significance alone.

To increase statistical power, we grouped events into composite breakage and amplification signatures and tested these against all three allelic TL measures. The breakage signature was specifically associated with shorter TL max, representing the longest allelic cluster on each arm. Arms harboring core breakage events (foldback inversions, foldback translocations, reciprocal inversions, and chromothripsis) had a TL max 0.98 kb shorter than unaffected arms (p = 0.019), and the full breakage signature showed a consistent effect (β = −0.68 kb, p = 0.005; Figure 4D). Breakage was not associated with TL min or TL median (all p > 0.05), though core breakage showed a marginal trend with TL median (β = −0.51 kb, p = 0.069). This allele-specificity is consistent with a detection model: while BFB cycles may initiate whenever a single telomere allele falls below a critical protective threshold, such events would remain subclonal and undetectable by bulk sequencing. Breakage events become detectable only when the arm’s telomeres are globally short (as reflected by a low TL max) indicating that a significant population of cellular lineages on that arm have crossed the dysfunction threshold.

The amplification signature showed the opposite pattern, associating specifically with longer TL min, or the shortest allelic cluster on each arm. Arms with core amplification events (multi-region intra-chromosomal ecDNA and duplicated inverted segments) had TL min 1.01 kb longer than unaffected arms (p = 0.004), and the full amplification signature showed a similar effect (β = +0.54 kb, p = 0.002; Figure 4E). Amplification was not associated with TL max (all p > 0.80). This indicates that amplification events occur preferentially on arms where even the shortest telomere allele in a population of cells is long, consistent with globally elongated telomeres maintained by ALT.

A systematic comparison of all signature and TL measure combinations confirmed the specificity and complementarity of these associations (Figure 4F). TL max was the only measure significantly associated with breakage signatures (core p = 0.019, full p = 0.005), while TL min showed the strongest associations with amplification signatures (core p = 0.004, full p = 0.002). TL median was also significantly associated with amplification (core p = 0.009, full p = 0.019) but not with breakage, showing intermediate sensitivity that reflects its position between the two allelic extremes. Together, these results support a dual-mode model in which chromosome-arm-specific telomere length governs two distinct pathways of structural genome evolution: arms with short telomeres are prone to breakage and loss, while arms with long ALT-maintained telomeres are prone to extrachromosomal amplification (Supplementary Figure 10).

After Benjamini-Hochberg correction for the 12 tests in the primary analysis (4 signatures x 3 TL measures), 6 associations remained significant at FDR < 0.05 (Supplementary Table 10). Both breakage associations with TL max survived correction (core q = 0.038; full q = 0.020), as did all amplification associations with TL min (core q = 0.020; full q = 0.019) and TL median (core q = 0.026; full q = 0.038). Leave-one-patient-out analysis confirmed that both primary associations were stable: the breakage core-TL max estimate was negative in 100% of iterations (90% individually significant at p < 0.05), and the amplification core-TL min estimate was positive in 100% of iterations (100% individually significant). Including tumor grade as a covariate did not materially alter the results (< 3% change in effect estimates), confirming that the dual-mode associations are not confounded by tumor progression stage.

### Structural variant breakpoints are enriched at telomeric and centromeric regions

Prior analyses established that arm-level telomere length is associated with two distinct modes of structural genome evolution: breakage on arms with short telomeres and amplification on arms with long telomeres (Figure 4). We next asked whether the physical positions of SV breakpoints are non-randomly distributed with respect to chromosome structure. The arm-specific concentration of inversions and BND-type variants observed in Figure 2G suggested that certain chromosomal regions may act as structural fragility hotspots. Here, we tested whether telomeric and centromeric compartments are preferentially affected.

To enable positional analysis across chromosomes of different sizes, we developed a genome-scaling framework that projects each breakpoint onto a normalized telomere–centromere–telomere coordinate axis, allowing direct comparison of breakpoint densities across arms and SV call sets (see Methods). Among SV classes, deletions and insertions showed the most consistent positional enrichment. In both the somatic (Severus) and germline-inclusive (Sniffles2) call sets, deletion and insertion breakpoints were recurrently enriched at telomeric and centromeric positions (Figure 5A; Supplementary Figure S12A), with chromosome-arm-wise deletion/insertion densities strongly correlated between the two call sets (Supplementary Figure S12B). Binning the scaled genome into telomeric, centromeric, and interstitial compartments confirmed this pattern quantitatively: deletions and insertion densities were highest at telomeres, intermediate at centromeres, and lowest in interstitial sequence in both somatic (Figure 5B) and total (Figure 5C) call sets (p < 0.001). Duplications showed a similar telomeric/centromeric bias, while inversions and BND-type variants exhibited more heterogeneous, arm-specific enrichment patterns (Supplementary Figure S12C), consistent with the focal hotspots identified previously (Supplementary Figure S2D).

**Figure 5.**
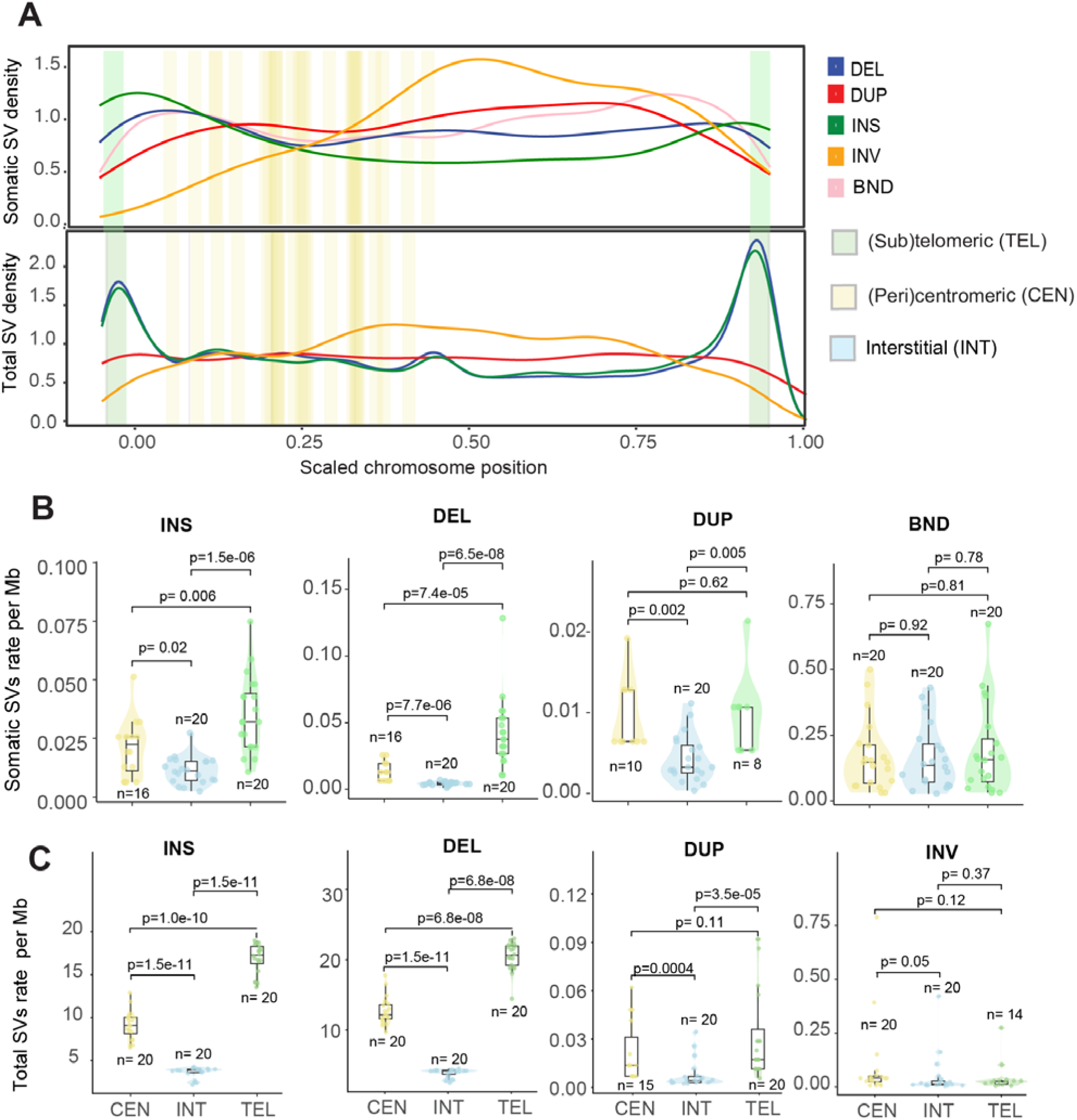
Structural variant breakpoints are enriched at telomeric and centromeric regions. **(A)** Genome-wide density profiles of SV breakpoints projected onto a normalized telomere–centromere–telomere coordinate axis across all non-acrocentric autosomal arms (n = 20 IDH-mutant astrocytoma tumor samples). Each line represents a chromosome, with genomic compartments shaded: telomeric/subtelomeric (green), interstitial (blue), and centromeric (khaki). **(B–C)** SV density (events per Mb) by genomic compartment for somatic SVs (B; Severus) and total SVs (C; Sniffles2). Distribution plots combine violin and boxplot geometries, with individual dots representing unique tumor samples to illustrate cohort-wide consistency. Bars represent mean density across samples. DEL and INS densities are significantly higher in telomeric and centromeric compartments than in interstitial regions (Kruskal-Wallis p < 0.001 for both call sets; pairwise comparisons by Wilcoxon rank-sum test with Bonferroni correction). See also Supplementary Figures S12–S13.

The concordance between somatic and germline-inclusive breakpoint landscapes is notable: it indicates that telomeric and centromeric compartments are constitutionally fragile and prone to structural variation regardless of whether events arise somatically or in the germline. This structural fragility likely reflects the repetitive sequence composition, late replication timing, and recombination-prone structure of subtelomeric and pericentromeric DNA, and operates independently of the ALT-driven, telomere-length-dependent breakage and amplification pathways described above. Importantly, these compartment-level trends were replicated in independent public glioma cohorts encompassing IDH-mutant astrocytomas, IDH-wild-type gliomas, and oligodendrogliomas (Supplementary Figure S13), indicating that telomeric/centromeric SV enrichment is a general feature of diffuse glioma genomes rather than a cohort-specific artifact.

Together, these findings identify two layers of structural vulnerability in IDH-mutant astrocytomas: a constitutive, heterochromatin-driven fragility at telomeric and centromeric regions that is shared across glioma subtypes and variant origins, and a telomere-length-dependent pathway in which ALT-driven heterogeneity in arm-specific telomere length shapes whether a given arm undergoes breakage or amplification.

## DISCUSSION

Our central finding is that chromosome-arm-specific telomere length governs two distinct modes of structural genome evolution in IDH-mutant astrocytoma. Arms on which even the longest telomere allele has fallen below a critical threshold preferentially harbor breakage-associated events (e.g., foldback inversions, chromothripsis, segmental losses) consistent with the classical model of telomere-dysfunction-driven BFB cycles (Maciejowski et al., 2020; Maciejowski et al., 2015; Umbreit et al., 2020). Conversely, arms on which even the shortest telomere allele remains long are enriched for ecDNA and complex amplification events, implicating ALT-mediated telomere elongation as a permissive or co-occurring feature of extrachromosomal amplification. The allele-specificity of these associations (TL max for breakage, TL min for amplification) is notable, because it reflects the population-level detection threshold in bulk sequencing: breakage becomes visible only when all subclonal lineages on an arm carry short telomeres, while amplification is detectable only when all lineages carry long telomeres. These results provide a model for understanding how ALT, by generating extreme inter-arm telomere length heterogeneity, simultaneously creates the conditions for both genome-destabilizing breakage and genome-amplifying ecDNA formation within the same tumor (Supplementary Figure 10).

The association between long telomeres and ecDNA raises a mechanistic question: how does ALT-mediated telomere elongation relate to extrachromosomal amplification? ALT is driven by break-induced replication (BIR) at telomeric ends, a recombination-based mechanism that is inherently mutagenic, involving long-range template switching, conservative DNA synthesis, and replication fork instability (Dilley et al., 2016; Zhang et al., 2019). While our genomic data cannot directly dissect mechanism, the co-occurrence of long telomeres and ecDNA on the same chromosome arms raises the possibility that ALT-associated recombination activity contributes to the chromosomal breakage and circularization events that generate ecDNA. In this model, the arms on which ALT is most active, reflected by globally long telomeres, may also be the arms most prone to BIR-associated rearrangements that produce extrachromosomal elements. Establishing a direct causal link will require carefully controlled functional experiments in model systems.

An alternative, well-established route to ecDNA biogenesis proceeds through chromothripsis and BFB cycles, in which chromosome shattering and subsequent reassembly can circularize amplified fragments (Shoshani et al., 2021). This pathway would be expected to operate on arms with critically short telomeres, where telomere dysfunction generates the dicentric chromosomes and chromatin bridges that trigger chromothripsis (Ly et al., 2019; Maciejowski et al., 2015). However, chromothripsis did not show the expected association with short telomeres in our cohort (univariate β = −0.39 kb, p = 0.890), despite contributing to the composite breakage signature. One possibility is that chromothripsis in ALT-driven tumors arises through a distinct mechanism, potentially through replicative processes such as microhomology-mediated BIR rather than classical NHEJ-dependent dicentric resolution (Cleal, Jones, Grimstead, Hendrickson, & Baird, 2019; Ngo et al., 2026). Alternatively, the small number of affected arms (n = 15) may simply have limited our statistical power. Resolving whether ALT tumors engage alternative routes to chromothripsis, and whether this contributes to ecDNA formation on long- or short-telomere arms, will require larger patient cohorts, higher resolution single-molecule long read approaches and careful experimental evaluation that can capture the temporal sequence of events and convincingly establish cause and effect.

Beyond the telomere-length-dependent pathways, our spatial enrichment analyses revealed a second, complementary layer of structural vulnerability. SV breakpoints were consistently enriched at telomeric and centromeric regions regardless of local telomere length, a pattern shared between somatic and germline-inclusive call sets and replicated across independent glioma cohorts including telomerase-positive glioblastomas and oligodendrogliomas. This constitutive fragility likely reflects the repetitive sequence composition and recombination-prone architecture of subtelomeric and pericentromeric DNA, and operates independently of ALT-driven telomere dynamics. These two layers, heterochromatin-driven positional fragility and telomere-length-dependent functional consequences, are not mutually exclusive. Rather, they suggest that certain chromosome arms are predisposed to structural variation by their sequence context, while ALT-driven telomere heterogeneity determines whether that variation manifests as breakage or amplification. The T2T-CHM13 reference genome, which resolves these previously inaccessible regions for the first time, was essential for this analysis (Nurk et al., 2022).

Several limitations should be acknowledged. First, the absence of matched germline sequencing makes it difficult to distinguish somatic from constitutional telomere length variation, and to definitively assign all SVs as somatic despite rigorous filtering. Second, read lengths in our dataset (mean N50 ∼12 kb) were shorter than currently achievable with optimized long-read protocols, constraining the accuracy of arm-level telomere length estimation by Telogator2, which requires reads to span into unique subtelomeric sequence for chromosome arm assignment. Future studies employing longer reads should substantially improve allele-specific telomere length resolution. We note that the core amplification signature was defined using multi-region intra-chromosomal ecDNA, a structurally complex subclass of ecDNA comprising multiple genomic fragments from the same chromosome, rather than all canonical ecDNA. This definition was chosen because multi-region intra-chromosomal ecDNA represents the most structurally elaborate circular elements and may more directly reflect the active BIR-mediated rearrangement processes underlying ecDNA biogenesis. Using all canonical ecDNA in the amplification signature yielded a non-significant association, suggesting that the telomere length relationship is specific to these complex elements rather than ecDNA as a broad category.

Third, the relatively small sample size (n = 20) limits statistical power for rare events such as chromothripsis and restricts generalizability. Fourth, the cross-sectional design cannot establish causality: we cannot determine whether short telomeres cause breakage or whether breakage erodes telomeres, nor whether long telomeres drive ecDNA formation or whether ALT preferentially elongates telomeres on arms already harboring ecDNA. Genomic analyses alone cannot functionally dissect mechanism; carefully controlled experimental studies will be essential to establish cause and effect. More broadly, many of the repeat-mediated structural variants detected in subtelomeric and pericentromeric regions involve ambiguous breakpoints that are difficult to represent in standard VCF format. Single-molecule long-read sequencing approaches capable of resolving complex rearrangements within these repetitive regions, combined with specialized variant representation frameworks, will be critical to fully characterize the structural consequences of telomere dysfunction and repeat-mediated instability. Longitudinal studies with matched primary-recurrent pairs and germline controls will further help disentangle these relationships.

In summary, our analyses establish the structural complexity of ALT-driven astrocytoma genomes and identify chromosome-arm-specific telomere length as a determinant of structural genome evolution. The dual-mode model, in which short telomeres drive breakage while long ALT-maintained telomeres co-occur with extrachromosomal amplification, provides a conceptual framework linking telomere biology to the diverse structural rearrangement patterns observed in these tumors, and motivates future investigation of BIR as a shared mechanism connecting ALT and ecDNA biogenesis.

## METHODS

### Patient cohort

Tumor samples were collected from *n* = 20 patients with histologically confirmed IDH-mutant astrocytomas (CNS WHO grades 2–3) treated at the Mayo Clinic. Key clinical characteristics, including age, sex and tumor grade are provided in Supplementary Table 1.

Genomic alterations were validated prior to long-read sequencing. The lack of 1p/19q codeletion status was confirmed using the OncoScan CNV Assay (Thermo Fisher Scientific) and Illumina EPIC array methylation according to the manufacturer’s protocol. Gene mutation status (including the presence of IDH mutation) was confirmed with directed sequencing with the Neuro-Oncology Expanded Gene Panel (Mayo Clinic Laboratories, test ID: NONCP).

All procedures were approved by the Mayo Clinic Institutional Review Board and performed in accordance with the Declaration of Helsinki. Written informed consent was obtained from all patients or their legally authorized representatives.

### Sample preparation and sequencing

Tumor tissue samples were obtained from patients with IDH-mutant astrocytomas undergoing surgical resection at Mayo Clinic, with approval from the institutional review board. Resected tissues were immediately transported to the pathology department for diagnostic evaluation and confirmation of tumor content. Samples selected for genomic analysis contained ≥70% tumor cells and were snap-frozen in liquid nitrogen within 30 minutes of resection. Tissues were stored at –80 °C until further processing.

Genomic DNA was extracted from 50–100 mg of frozen tissue using the QIAamp DNA Mini Kit (Qiagen) or the AllPrep DNA Mini Kit (Qiagen), according to the manufacturer’s instructions. Tissue homogenization was performed using a bead mill homogenizer to ensure complete lysis. DNA concentration was quantified using the Qubit dsDNA BR Assay Kit (Thermo Fisher Scientific), and purity was assessed by NanoDrop spectrophotometry (Thermo Fisher Scientific), with acceptable A260/280 ratios of ∼1.8. DNA integrity was evaluated using pulsed-field gel electrophoresis or the Agilent 4200 TapeStation system. DNA samples with high molecular weight and adequate purity were used for downstream genomic analyses.

Library preparation was performed using Oxford Nanopore Technologies (ONT) Ligation Sequencing Kit (SQK-LSK114). DNA was sheared to ∼20–30 kb using g-TUBEs (Covaris). End-repair, A-tailing, barcode ligation, and sequencing adapter attachment were done per ONT protocol. Sequencing was performed on R10.4.1 flow cells (FLO-PRO114M) using a PromethION device. For four patient samples sequenced across two flow cells, libraries from both runs were merged to increase coverage (>15×), ensuring sufficient depth for downstream analyses.

### Basecalling and adapter trimming

Raw signal data were basecalled using Dorado v7.1.4 with the high-accuracy model dna_r10.4.1_e8.2_400bps_hac@v4.2.0. Adapter sequences were automatically trimmed during basecalling. Basecalled reads were converted into unaligned BAM (uBAM) files, which were subsequently converted to FASTQ format using Samtools v1.21 (H. Li et al., 2009). Read quality was assessed using NanoPlot, and low-quality or chimeric reads (Q < 7, length <1 kb) were filtered using NanoFilt (De Coster & Rademakers, 2023). Only high-quality reads passing these filters were carried forward for alignment.

### Genome alignment

High-quality reads were aligned to the human T2T-CHM13 v2.0 reference genome using Minimap2 v2.24 (Li, 2018) with the map-ont preset optimized for Oxford Nanopore reads. For four samples sequenced across two flowcells, reads from both runs were merged prior to alignment. The resulting BAM files were sorted and indexed using samtools v1.21 (Heng Li et al., 2009), and alignment quality was assessed using stats. Only properly aligned primary reads were retained for downstream SV and CNV analyses.

### SVs calling

We developed a custom pipeline to process BAM files and generate both germline-inclusive and somatic variant calls, allowing flexible selection of analysis modes (GitHub: https://github.com/barthel-lab/LongSV_caller). The pipeline integrates multiple SV callers: Sniffles2 v2.7.1 for germline-inclusive variants (with optional “--mosaic” mode), Severus v0.1.2 for somatic SV detection in tumor/normal or tumor-only settings. All tools are orchestrated through a Snakemake workflow, enabling reproducible and scalable execution across datasets.

In this study, the custom SV detection pipeline was executed on all samples, processing each tumor BAM individually. For downstream analyses, we specifically focused on germline-inclusive SVs called by Sniffles2 and somatic SVs identified by Severus, enabling a consistent and high-confidence assessment of structural variation across the cohort. Severus was run using the CHM13 VNTR BED file to account for tandem repeat regions (--vntr-bed) and a panel of normals (--PON) to enhance the accuracy of somatic variant detection. This setup enables accurate detection of tumor-specific SVs without requiring a matched normal sample.

### Haplotype-aware copy-number profiling

To achieve high-resolution haplotype-aware reconstruction, we first called small variants and performed haplotype phasing using Clair3 (Zheng et al., 2022) together with Longphase (Lin, Chen, Yu, & Huang, 2022). Phased BAMs were then analyzed with Wakhan v0.4.2 (Ahmad et al., 2025); GitHub repository: KolmogorovLab/Wakhan) for haplotype correction, purity/ploidy estimation, and haplotype-specific copy-number profiling. Wakhan analyses incorporated SV breakpoint information from Severus to refine copy-number segment boundaries and support downstream interpretation complex rearrangement structure.

### SVs refinement, manual curation, and downstream processing

To refine somatic SV detection, Severus was run in multi-sample mode to identify shared events across the cohort, which were used to build a customized panel of normals (PoN). SVs observed in at least two samples were considered likely technical or germline artifacts. These recurrent events were combined with the CHM13-based PoN (Schloissnig et al., 2025) provided in the Severus repository to generate an expanded PoN for downstream filtering. Each tumor BAM was then reprocessed in single-sample mode using the updated PoN to prioritize tumor-specific SVs. To further reduce residual germline contamination, we applied an allele-fraction filter using per-event VAF and haplotype-resolved VAF (hVAF) when available; most calls clustered at low VAF/hVAF, whereas a subset showed hVAF ≈ 1. We retained SVs with hVAF ≤ 0.8 and, for unphased regions, VAF ≤ 0.4, excluding events above these thresholds as likely germline. Final per-sample VCFs were compressed and indexed with bgzip and tabix. Individual filtered VCFs were then merged into a multi-sample VCF using bcftools merge (v1.17) (Danecek et al., 2021). The merged VCF was converted to BED format using bcftools query to extract chromosomal coordinates, SV types, and sample identifiers for integrative analyses with telomere length and other genomic features.

Complex SV clusters identified by Severus were manually annotated using multiple signals, including the DETAILED_TYPE annotation in the VCF, BND profiles, and haplotype-resolved copy-number states derived from Wakhan. Classification was guided by established patterns of rearrangement topology and copy-number context. Clusters were assigned to the following categories: chromothripsis-like events, chromoplexy-like events, templated insertion cycles/ chains, foldback inversion–associated rearrangements, and other complex events spanning either single chromosome arms or multiple chromosomes.

For downstream analyses, germline-inclusive and somatic SVs were analyzed separately. All analyses were performed relative to the CHM13 human reference genome, which provides a complete assembly of previously unresolved regions, including centromeres and telomeres.

### CNV profiling from long-read sequencing

To assess CNVs across the cohort, we developed an alignment-free, k-mer-based coverage estimation approach to mitigate sequence composition bias and avoid multi-mapping artifacts in repetitive regions. For each sample, a k-mer database (k = 31) was constructed from sequencing reads using KMC (Kokot, Dlugosz, & Deorowicz, 2017). Each database was then intersected with a precomputed set of reference-unique 31-mers from the T2T-CHM13 v2.0 assembly, retaining only k-mers that occur exactly once in the reference genome. This restriction to unique k-mers eliminates coverage inflation from repetitive and multi-copy sequences, effectively normalizing for sequence composition without requiring explicit GC-correction. Per-position coverage of unique k-mers along each chromosome was computed using a custom lookup tool, and coverage values were aggregated into 200 kb genomic bins by computing the mean coverage of all unique k-mers within each bin. Per-sample binned coverage profiles were then merged into a single cohort-wide matrix for downstream segmentation.

To eliminate artifactual signals in read-depth coverage, genomic regions overlapping blacklist regions; such as centromeres, telomeres, and sex chromosomes (chrX and chrY); were excluded using Bedtools v2.30.0 (Quinlan & Hall, 2010) with the intersect -v command. This filtering step minimized false positives arising from low-complexity and structurally variable genomic regions.

Next, log ratio values were computed for each sample by dividing its normalized coverage by the median reference coverage across all samples, followed by log transformation (Zhao et al., 2004). Sample names were standardized and the data reshaped to matrix format using tidyverse tools in R.

Segmentation of the log ratio data was performed using the DNAcopy v1.85.0 R package. Circular Binary Segmentation (CBS) algorithm (Olshen et al., 2004) was applied to identify regions of significant copy number gains or losses. A CNA object was created using the log - transformed coverage matrix, chromosomal positions, and sample identifiers. The final segmented CNVs were exported in BED format for integration with structural variation and telomere length data. CNVs were categorized according to segment mean values: segments with seg. mean ≥ 0.2 were classified as gains, seg. mean ≤ -0.2 as losses, and all others as neutral.

DNAcopy-derived segments were used for standardized gain and loss burden analyses across samples and to test for telomere length associations. Wakhan was added at a later point for the haplotype-aware copy-number reconstruction and for the interpretation of complex structural rearrangements.

### Detection and characterization of ecDNA and High Gains (HGs)

Extrachromosomal DNA (ecDNA) was reconstructed from long-read sequencing data using Decoil v1.1.2-slim (Giurgiu et al., 2024) within a Singularity container (Decoil.sif) to ensure a reproducible analysis environment. This tool integrates structural variant breakpoints with sequencing coverage to reconstruct the complete topology of each element and estimate its proportional abundance relative to the linear genome. To ensure high-confidence reconstructions, Decoil employs a LASSO regression model to prune the candidate cycle set, removing reconstructed paths with estimated proportions below the mean whole-genome sequencing (WGS) coverage (Giurgiu et al., 2024). Input files consisted of coordinate-sorted and indexed BAMs derived from long-read minimap alignments. The pipeline generates per-sample outputs, including ecDNA loci, fragment topology (simple vs. complex), and estimated relative abundances, allowing accurate characterization of tumor-specific circular DNA structures. Following these outputs, circular elements >0.1 Mb were classified as canonical ecDNA, while smaller fragments were categorized as short circular DNAs (Giurgiu et al., 2024). Decoil further assigns each reconstruction to one of its seven hierarchical topologies of increasing complexity, ranging from simple circularization to complex foldback inversions (Giurgiu et al., 2024). Based on these high-confidence outputs, we calculated ecDNA burden for each chromosome arm as the arm-level ecDNA proportion (estimated_proportions), which quantifies the fraction of total ecDNA signal assigned to a given arm. With the exception of sample 6436B, which was omitted due to an unexpected pipeline failure, all samples were successfully processed and included in the final arm-level calculations.

Visualization of ecDNA structures was performed using Decoil’s Decoil-viz module within the same Singularity environment. This produced interactive HTML reports and organized output folders summarizing fragment structures, genomic coordinates, and relative abundances per sample. These visualizations enabled assessment of fragment topology, circularization patterns, and sample-level ecDNA burden.

Decoil reports each reconstructed circular element as a set of one or more genomic fragments that collectively define the circular topology. We report element-level counts (unique Decoil circ_ids from the summary output) rather than fragment-level counts, as a single ecDNA circle may comprise multiple genomic segments. Elements labeled "ecDNA" by Decoil were classified as canonical ecDNA. All remaining elements were classified as short circular DNAs.

In parallel, highly gained regions (HGs) were identified using the median absolute deviation (MAD). For each sample, CNV segment means were used to calculate the median and MAD, and segments exceeding the median by ≥4× MAD with lengths between 10 kb and 5 Mb were classified as HGs, representing outlier gain events relative to the sample background. HGs coordinates were further annotated against CHM13 exons and repetitive elements, including LINEs, SINEs, and Alu sequences, to characterize their sequence composition.

### Arm-level aneuploidy analysis

Arm-level aneuploidy was quantified using an in-house R pipeline, conceptually following the approach described for the TCGA pan-cancer analysis (Taylor et al., 2018). Briefly, segmented copy number profiles were used to calculate discrete copy number alterations for each chromosomal arm. Gains and losses were defined based on deviation thresholds from sample-specific baseline ploidy. For each sample, an arm-level aneuploidy score was computed by summing the total number of gained and lost arms across the genome, providing a quantitative measure of chromosomal imbalance.

These scores were used to assess global patterns of aneuploidy and were integrated with structural variant and telomere length data for comparative analyses across samples.

### Arm-level SV/CNVs normalization and heterogeneity

To accurately assess and compare the distribution of SVs and CNVs across the genome, burden calculations were normalized by genomic scale. For each sample, the number and cumulative size of each variant type (e.g., deletions, duplications, insertions, inversions for SVs; gains and losses for CNVs) were computed per chromosome arm (p and q). Variant coordinates were assigned to chromosome arms using a cytoband annotation file adapted to the CHM13 v2.0 reference genome, which defines centromere and telomere positions as well as arm boundaries. Chromosome and arm lengths were derived from this annotation to enable normalization of variant burden by genomic interval size (Kent et al., 2002).

We quantified per-arm variant burden as an event rate per megabase (events/Mb) by dividing the number of variants on each chromosome arm by its length. Arm-level burdens were then normalized by total chromosome length to reduce chromosome-wide size effects and support proportional comparisons across samples and chromosomal regions. This arm-resolved, length-normalized framework allowed precise quantification of structural alteration burdens per genomic unit. We summarized arm-level heterogeneity using two complementary measures. First, we computed the Gini coefficient using the ineq library in R to quantify how unevenly SV burden is distributed across chromosome arms (higher values indicate greater concentration in a subset of arms). Second, we reported prevalence (P) as the fraction of chromosome arms with non-zero SV burden (x_i_ > 0), capturing the genome-wide spread of events. For SV burdens *x*_1_ ≤ ... ≤*x_n_* across *n* arms (sorted in ascending order) with mean *µ*, the Gini index was computed as:

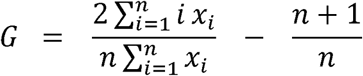

Here, *G* = 0, indicates a uniform SV burden across arms, whereas larger values indicate increasing concentration of burden within a subset of arms (hotspot-driven heterogeneity). Visualization and clustering were performed using the ComplexHeatmap (Gu, Eils, & Schlesner, 2016) R package and Python’s seaborn (v0.13.2) and matplotlib libraries (Hunter, 2007)

### Telomere length estimation

Allele-specific telomere length (TL) was estimated from Oxford Nanopore Technology (ONT) long-read whole-genome sequencing data using Telogator2 (Stephens et al., 2022; Stephens & Kocher, 2024). Telogator2 identifies telomere-containing reads, clusters them by telomere variant repeat (TVR) composition to resolve individual alleles, and anchors each allele to a chromosome arm by aligning the subtelomeric portion of the read against a panel of T2T reference assemblies (T2T-CHM13, T2T-HG002, T2T-YAO, T2T-CN1, T2T-I002C; both maternal and paternal haplotypes where available). Alleles flagged as interstitial telomeres were excluded from all downstream analyses (n=60 reads removed). For each sample and chromosome arm, three summary statistics were derived from the per-allele TL estimates (75th percentile method, TL_p75): the minimum (TL Min, shortest allele), maximum (TL Max, longest allele), and median (TL Median) across all alleles assigned to that arm. Because Telogator2’s TVR-based clustering can resolve more allele clusters than the true sample ploidy when reads anchor to different reference builds, min/max/median aggregation across alleles was used rather than selecting a single representative allele per arm.

### Telomere length normalization and categorization

For cross-sample comparison and visualization, arm-level TL Median values were globally z-score normalized by subtracting the cohort-wide mean (4.97 kb) and dividing by the cohort-wide standard deviation (1.80 kb) across all 780 sample-arm observations. This global z-score preserves both between-patient and between-arm TL differences, enabling identification of samples with globally short or long telomeres. Chromosome arms were categorized based on global z-score thresholds: critically short (z < -1.5), short (-1.5 <= z < -0.5), normal (-0.5 <= z < 0.5), long (0.5 <= z < 1.5), and very long (z >= 1.5).

### Genomic event signatures

Chromosome arm-level genomic events were classified into two composite signatures:

1. **Breakage signature.** Events associated with telomere-driven chromosome breakage and loss. The core breakage signature scored an arm as positive if it harbored any of the following: foldback inversions detected by Severus breakpoint clustering (auto_foldback_Count > 0), foldback translocations identified through manual review of Wakhan haplotype-resolved reconstructions (Foldback_Translocation > 0), reciprocal inversions called by Severus (sv_count_INV > 0), or chromothripsis identified through manual Wakhan review (Chromothripsis > 0). The core breakage signature uses the VCF-level reciprocal inversion count. The full breakage signature additionally included arms with any segmental copy number loss (cnv_rate_per_mb_Loss > 0) or arm-level aneuploidy loss.
2. **Amplification signature.** Events associated with telomere-mediated amplification and extrachromosomal DNA (ecDNA) formation. The core amplification signature included arms harboring multi-region intra-chromosomal ecDNA reconstructed by Decoil (ecdna_multi_region_intra_chr > 0) or duplicated inverted segments detected by Severus (auto_dup_inv_segment_Count > 0). The full amplification signature additionally included arms with HG or ecDNA (ecDNA_count > 0), complex inversions (auto_complex_inv_Count > 0), or single- and multi-chromosome complex rearrangements.

Each chromosome arm was scored as positive (>=1 event) or negative (0 events) for each signature.

### Statistical analysis

Associations between telomere length and genomic event signatures were tested using linear mixed-effects models with the form TL ∼ signature + (1|Sample_ID), where Sample_ID was included as a random intercept to account for within-patient correlation across chromosome arms (∼39 arms per sample). All arm-level analyses were restricted to non-acrocentric autosomal arms. For the sample-level comparison of tumors with versus without HG/ecDNA events, the per-sample median TL was computed as the median of arm-level TL Median values across all chromosome arms, and groups were compared using the Wilcoxon rank-sum test. Global z-score permutation tests (10,000 permutations) were used to assess whether arms harboring specific event types had significantly shifted TL distributions compared to the cohort background. All analyses included 20 IDH-mutant astrocytoma samples comprising 745 sample-arm observations. The association between chromothripsis and tumor grade was assessed using Fisher’s exact test on the sample-level 2×2 contingency table (chromothripsis present/absent × grade 2/3), which is preferred over the chi-squared test when expected cell counts are below 5.

To account for the multiple hypothesis testing burden in the primary signature analysis, Benjamini–Hochberg false discovery rate (FDR) correction was applied across the 12 tests in the composite signature × TL measure matrix (4 signatures × 3 TL measures). Corrected q-values are reported alongside nominal p-values (Supplementary Table 10). As additional robustness checks, leave-one-patient-out analysis was performed by refitting each primary model 20 times with one patient excluded per iteration to assess stability, and tumor grade was included as a fixed-effect covariate to test for confounding by tumor progression stage.

### Unsupervised clustering of telomere length profiles

To identify patient subgroups based on telomere length profiles, the median allelic TL per chromosome arm was organized into a 20-sample x 39-arm matrix and globally z-score normalized. Hierarchical clustering was performed on both samples (rows) and chromosome arms (columns) using Ward’s linkage on Euclidean distances with optimal leaf ordering. Sample-arm pairs lacking Telogator2 measurements (n=43 of 780, 5.5%) were imputed with the per-arm median prior to clustering. Row annotations included tumor grade (available for all 20 samples) and presence/absence of HG/ecDNA events.

### Spatial enrichment analysis of SV in subtelomeric and pericentromeric regions

To explore the spatial distribution of somatic and germline-inclusive SVs relative to chromosomal hinges—namely subtelomeres and pericentromeres, we calculated the distance of each SV and CNV to key chromosomal landmarks using the CHM13 reference genome. Centromere positions were precisely defined based on CHM13 annotations, with flanking regions extending ±50 kb (0.05 Mb) on both arms designated as pericentromeric zones. Telomere ends were defined separately for each chromosome arm using curated telomeric coordinates from the same reference. Subtelomeric regions were delineated as the terminal 3% of the chromosome length on each arm, corresponding to scaled genomic positions <0.03 for the p arm and >0.97 for the q arm, based on normalized chromosome coordinates.

To enable direct comparison of variant positions across chromosomes of different lengths, we rescaled genomic coordinates to a normalized 0–1 axis for each chromosome. For each variant, we defined the breakpoint midpoint as midpoint = (start + end)/2 and normalized it to the chromosome span from the start of the p arm (start_*p*_) to the end of the q arm (end_*q*_):

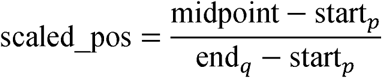

The centromere was mapped onto the same scale using the centromere start coordinate:

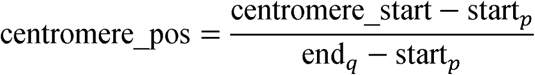

This normalization places all variants in a common coordinate system, allowing cross-chromosomal comparisons of positional enrichment near telomeric, centromeric, and interstitial regions. For downstream analyses, each chromosome was partitioned into p and q arms at the normalized centromere position. To visualize the spatial distribution and burden of genomic alterations, we evaluated SV density across the genome using line plots and density plots, with additional normalization per megabase (Mb) to adjust for chromosome size. These visualizations revealed local enrichments and depletions across different genomic zones. Furthermore, we statistically compared the variant distributions between hinge regions (subtelomeric and pericentromeric) and interstitial genomic regions using Wilcoxon rank-sum tests to assess significant enrichment or depletion of variants near chromosome ends and centromeres.

### Public dataset validation of glioma-associated variant and telomere signatures

To validate SV and CNV signatures alongside telomere profiles observed in our cohort, we re-analyzed publicly available datasets from the The Cancer Genome Atlas Lower Grade Glioma (TCGA-LGG) and Glioma Longitudinal AnalySiS (GLASS) Consortium glioma cohorts (Ceccarelli et al., 2016; Varn et al., 2022). These analyses included IDH-mutant/1p19q-codeleted, IDH-mutant/1p19q-intact, and IDH-wildtype gliomas, with samples further categorized as primary or recurrent tumors where information was available. In total, the cohort comprised 760 samples, including 430 primary and 330 recurrent tumors. From this dataset, we identified a high-confidence longitudinal cohort of 272 paired patients (544 samples total) with matching primary and recurrent copy number calls. Telomere length estimates derived from whole-genome sequencing were integrated with structural variant and copy number variation data accessed via cBioPortal and the GLASS data portals.

For CNV burden classification, segmentation files were used to distinguish aneuploid events (chromosome arms with ≥80% gain or loss) from segmental CNVs (all other events). This enabled quantification of arm-level and segmental CNV burdens and assessment of aneuploidy across IDH subtypes and recurrent tumors. By applying the same analytical pipelines for SV and CNV characterization, we examined how genomic instability and telomere attrition co-occur across samples. Cross-cohort comparisons confirmed the robustness of these associations and underscored the biological relevance of variant burden and telomere dynamics in glioma progression.

## ACKNOWLEDGEMENTS

This work was supported by the Department of Defense (award HT9425-23-1-0844 to F.P.B.), the Lennar Foundation and the National Cancer Institute (R01CA298083 to J.E-P. and R01CA230712 to R.B.J). Oxford Nanopore Technologies, together with the Mayo Clinic Advanced Diagnostics Laboratory in the Department of Laboratory Medicine and Pathology supported the long read PromethION sequencing. Large language models were used in the preparation of this manuscript to improve grammatical accuracy and readability. The authors reviewed all edits and retain sole responsibility for the accuracy and integrity of the content. We gratefully acknowledge the Mayo Clinic Neuro-Oncology Biorepository and the patients and families whose participation in research made this study possible.

## DATA AND CODE AVAILABILITY

Datasets described here can be found on Synapse at https://www.synapse.org/Synapse:syn72016012). The code used in this analysis is available from GitHub (https://github.com/barthel-lab/telomere-SV-analysis).

