## Supplementary Figures for "Chromosome-arm-specific telomere length governs dual modes of structural genome evolution in IDH-mutant astrocytoma"

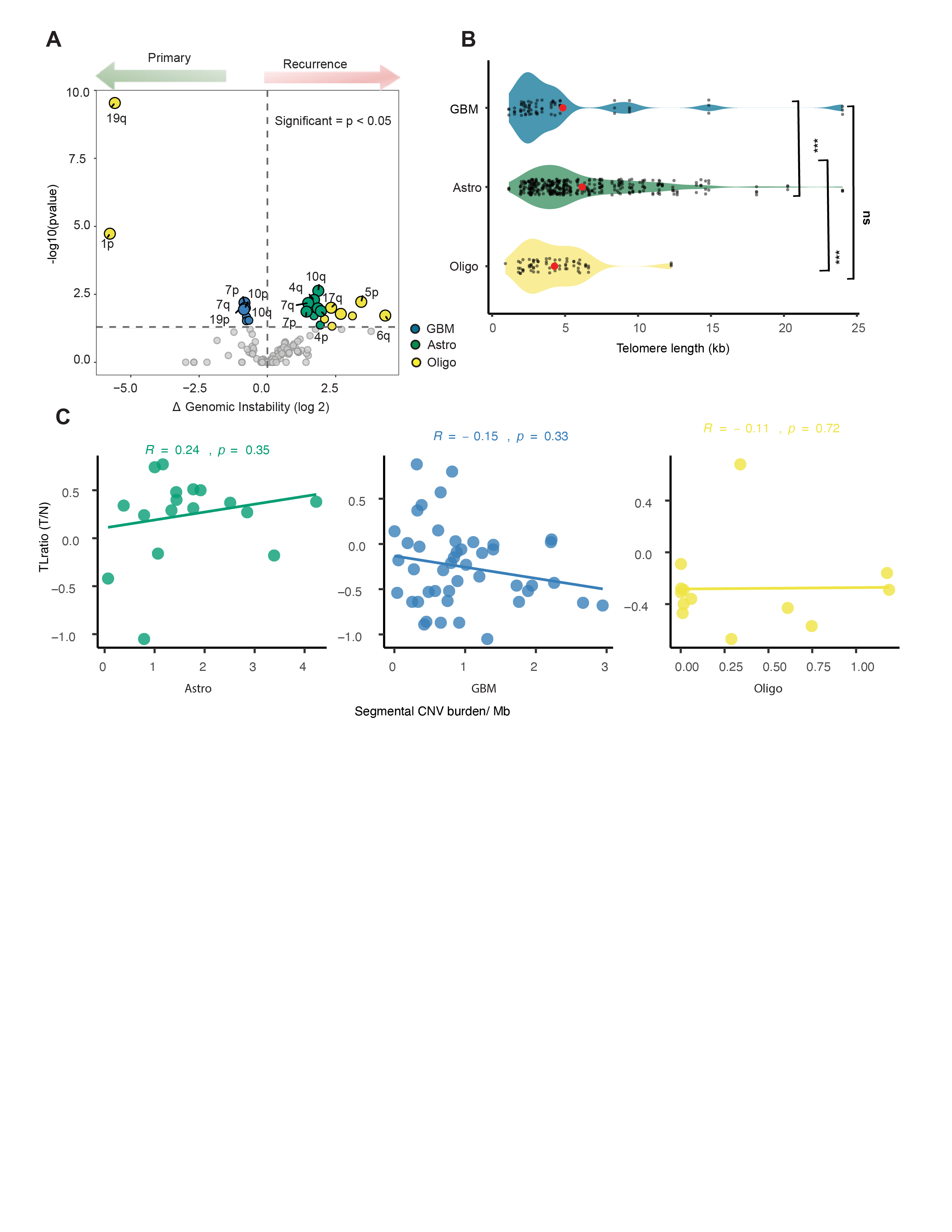

**Supplementary Figure 1. Telomere dynamics and genome instability across diffuse glioma subtypes** (A) Volcano plot displaying differential genome instability between primary and recurrent tumors, highlighting chromosome arms with significant CNV enrichment in recurrent samples (adjusted P < 0.05, Wilcoxon rank-sum test). Arms with positive log2 fold change denote increased instability upon recurrence. (B) Violin plots showing telomere length distributions across glioma subtypes, derived from long-read sequencing. Astrocytoma displayed significantly longer and more heterogeneous telomeres relative to oligodendrogliomas and glioblastomas (P < 0.0001, Wilcoxon test), consistent with enhanced telomere erosion and ALT-associated telomere maintenance. (C) Spearman correlation analyses reveal divergent subtype-specific associations between telomere T/N ratio and segmental CNVs burden across astrocytomas, glioblastomas, and oligodendrogliomas.

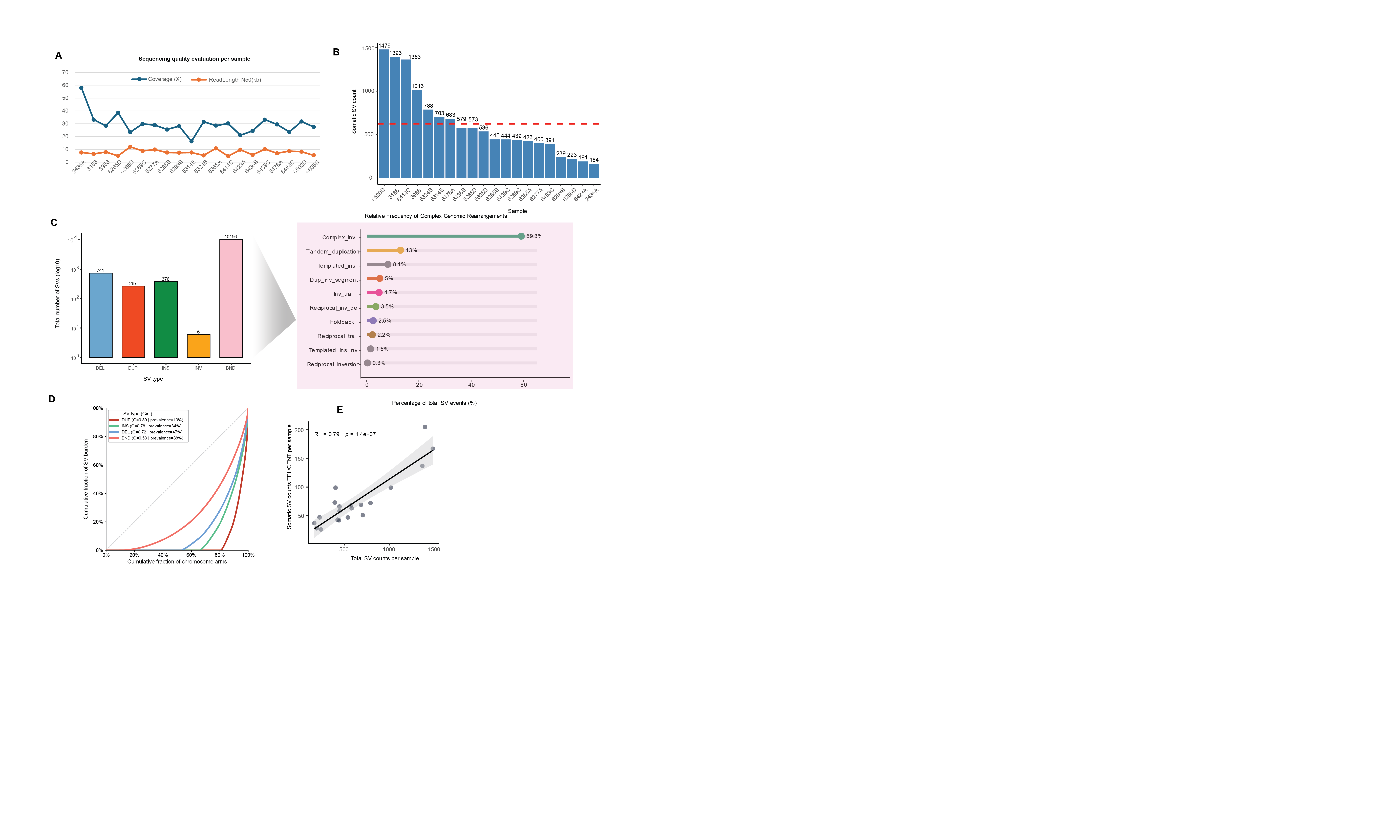

**Supplementary Figure 2. Somatic structural variant burden per sample and chromosome arms. (A)** Line plots showing read length N50 (orange bars) and sequencing coverage (blue bars) for each sample on the x-axis. The y-axis represents both metrics with appropriate scaling, illustrating variation in read length distribution (N50) and sequencing depth (coverage) across samples in a single comparative plot. Bar plots summarize the SV landscape across all tumors. **(B)** *(Left)* Total SV counts per sample, with the red dotted line indicating the cohort-wide mean. This plot highlights substantial inter-tumoral variability in overall SV load. *(Right)* Breakdown of complex genomic rearrangements: this plot illustrates the relative frequency of complex rearrangement types across the cohort. Percentages and raw counts are annotated to highlight the predominance of specific SV signatures **(C)** SV counts classified by SV class, with distinct colors representing insertions, deletions, duplications, inversions, and BND. BNDs are the most abundant SV class across the cohort, occurring at significantly higher frequencies than other SV types, reflecting extensive breakpoint fragmentation in these samples. **(D) Lorenz curves** depict the cumulative fraction of total SV burden (y-axis) explained by the cumulative fraction of chromosome arms (x-axis) for each SV type (arm-level median burden aggregated across samples). The dashed diagonal indicates perfect equality (uniform burden across arms). Increased curvature indicates stronger concentration of SV burden within a subset of arms (hotspot-driven instability). The on-plot annotation reports the **Gini index (G)** (0 = uniform, 1 = highly concentrated) and **prevalence (P)**, defined as the percentage of chromosome arms with non-zero SV burden (arm-level median burden > 0).
**(E)** Scatter plot shows, for each sample, the total SV count (x-axis) versus the number of telomeric/centromeric SVs (y-axis), with SVs called and annotated against the CHM13 reference genome. Association strength and significance were assessed using a Spearman correlation test.

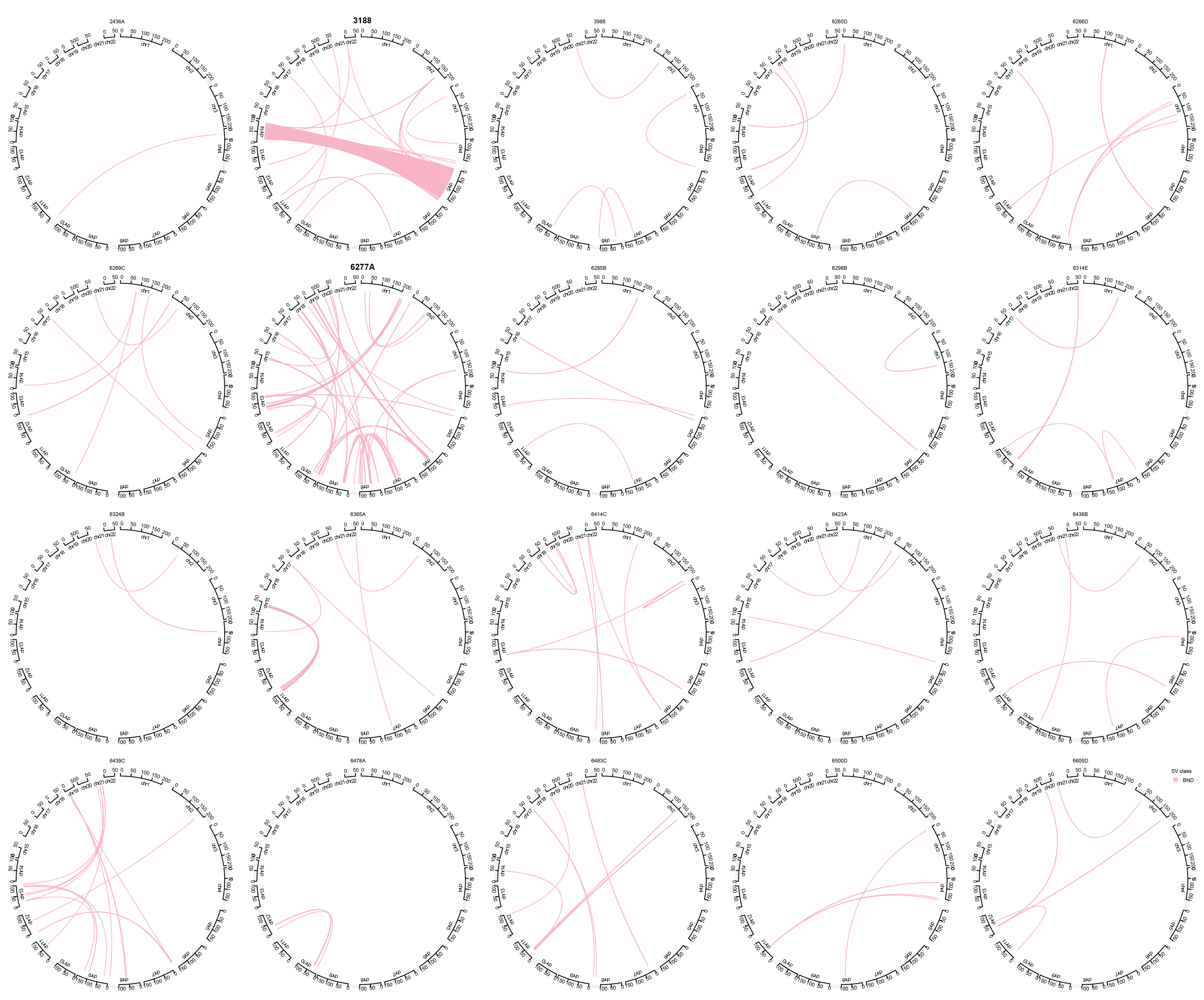

**Supplementary Figure 3. Genome-wide mapping of interchromosomal breakpoints (BND) events using Circos visualization.** Circos plots illustrate the full spectrum of **BND-mediated interchromosomal rearrangements** detected in each astrocytoma sample. The **outermost ring** represents the T2T-CHM13 chromosome ideograms arranged in genomic order, providing reference coordinates for all breakpoint positions. **Pink arcs** denote individual BND events, each linking two genomic loci involved in an interchromosomal translocation; the arc density reflects the degree of large-scale genome fragmentation within each tumor. Samples **3188** and **6277** show **markedly elevated translocation burden**, characterized by dense radial networks of arcs spanning many chromosomes, consistent with **extensive chromosomal shattering.**

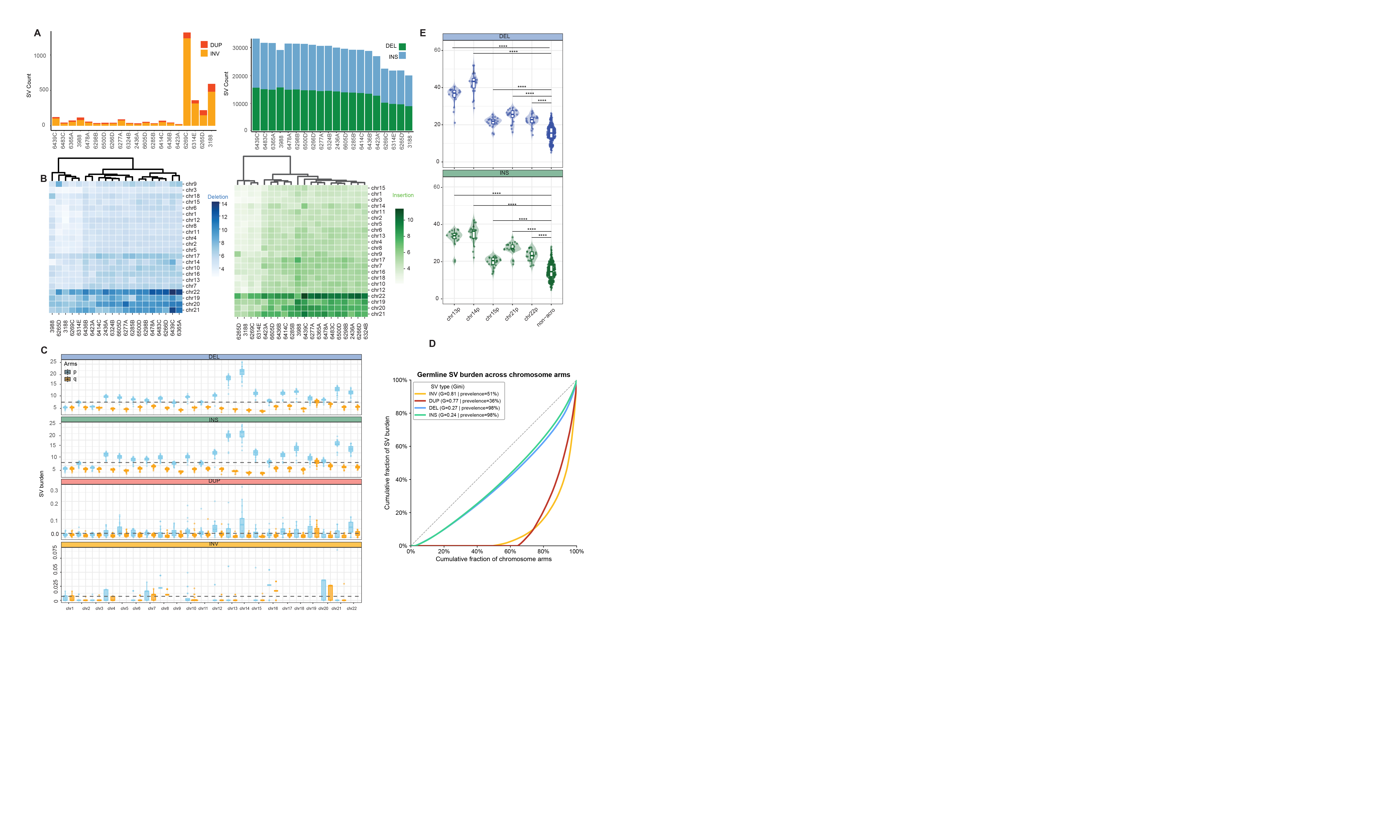

**Supplementary Figure 4. Germline structural variant summary across samples and chromosome arms.** (**A)** SV ratio of different types of structural variants across clinical samples, indicating abundance of insertions and deletions in astrocytomas cohorts. (**B)** Chromosome-wide heatmaps displaying genomic SVs identified through long-read sequencing. The analysis highlights a high abundance of insertions (green) and deletions (blue), which are particularly enriched in shorter chromosomes, as evident from the clustering pattern. **(C**) SV burden normalized by megabases (Mb) for each chromosome arm. The x-axis displays chromosomes separated into p and q arms, with p arms highlighted in orange and q arms in sky blue. Above each arm, grouped bars represent SV burdens colored by SV type: DEL in blue, DUP in red, INV in orange, and INS in green. Notably, the acrocentric p arms of chromosomes 14, 21, and 22 show elevated insertion and deletion burdens compared to other chromosome arms. Significant differences observed in SV burdens confirm distinct mutational patterns associated with acrocentric chromosome arms **(D)** Lorenz curves and Gini indices representing the cohort-wide distribution of germline SVs. The near-linear profile and low coefficients demonstrate a systemic, uniform distribution that contrasts with the focal landscape of somatic variation. **(E)** Statistical comparison of SV burdens between acrocentric and non-acrocentric chromosome arms for INS and DEL SV type events, showing increased INS and DEL burdens in acrocentric arms. SV burden was quantified at the chromosome-arm level, and differences between groups were assessed using a two-sided Wilcoxon rank-sum test, with significance denoted by asterisks, are indicated in the plot.

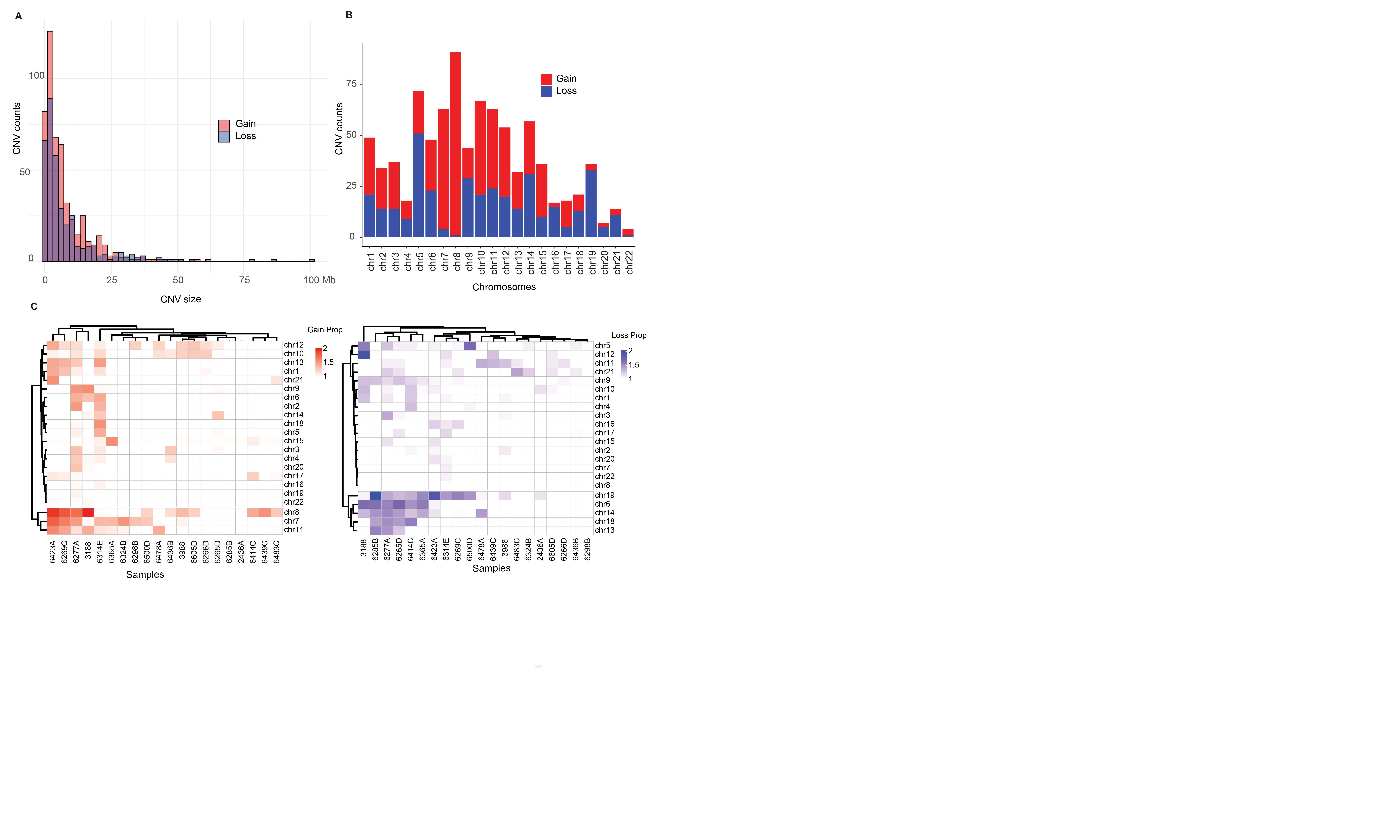

**Supplementary Figure 5. Copy Number Variation (CNV) Patterns and Clustering Across the Cohort**. **(A)** Histogram showing the distribution of CNV event sizes, separated into **gains** and **losses**. Gains are more frequent and generally smaller in size, while losses are less frequent but span larger genomic regions. **(B)** Bar plot showing the total number of CNV gains and losses per chromosome across the cohort. The y-axis represents CNV count, and the x-axis lists chromosomes. This reveals chromosome-specific CNV burden patterns, with some chromosomes more frequently altered. **(C)** Heatmaps depicting the distribution of CNV gains and losses across all samples. Rows represent chromosomes and columns represent individual samples. Color intensity indicates the relative proportion per MB of CNV events (gain or loss). Hierarchical clustering using a dendrogram was applied to both samples and chromosomes based on CNV proportions, revealing distinct clusters of samples and chromosomal regions with recurrent copy number alterations.

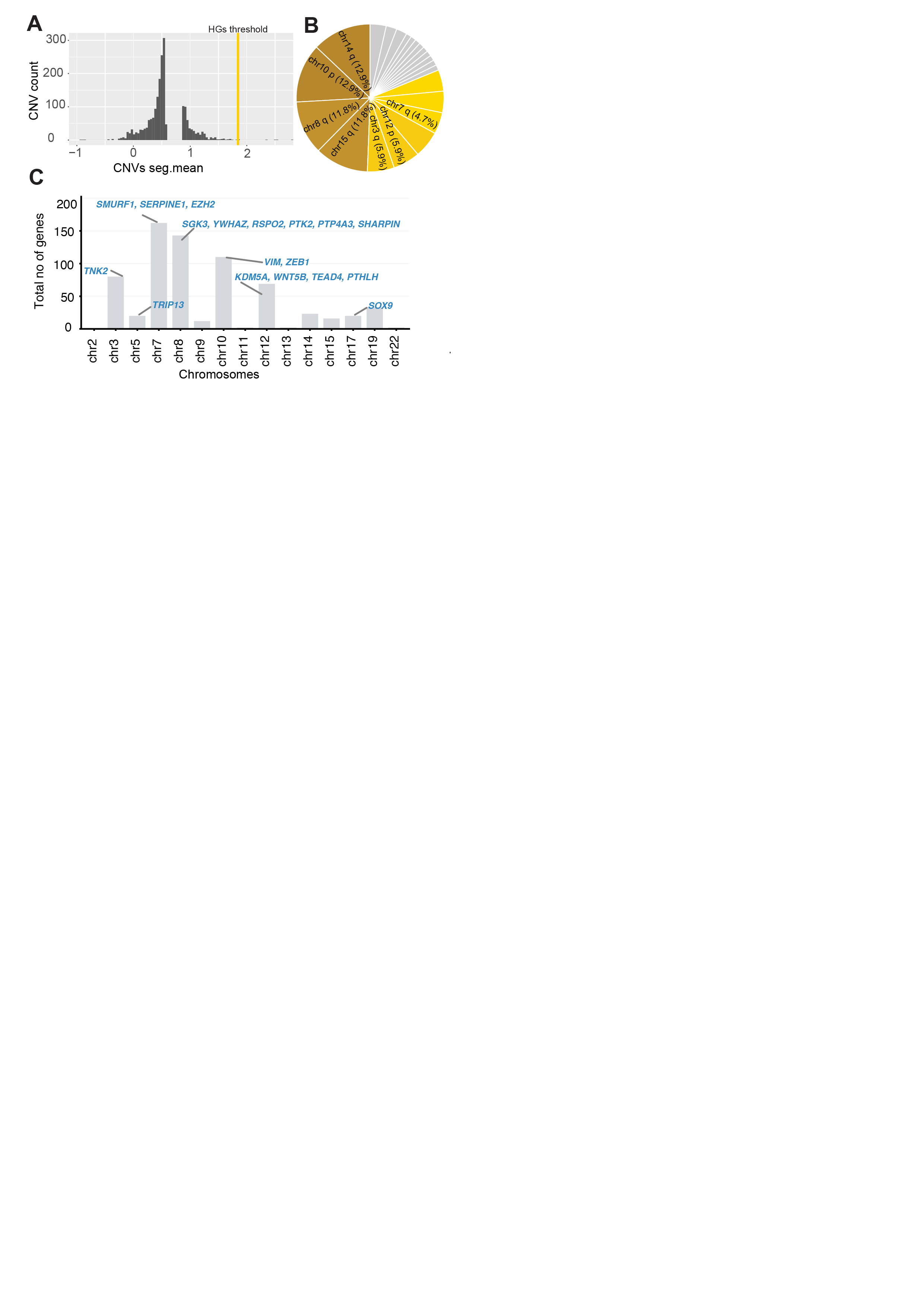

**Supplementary Figure 6. Genome-wide detection and functional characterization of High Gains (HGs) in diffuse glioma.** **(A)** Histogram of segmented copy-number means across all samples. The yellow vertical line denotes the literature-supported amplification threshold that defines High Gains (HGs), distinguishing high focal amplifications from baseline CNV variation. **(B)** Circular plot showing the proportion of chromosome arms harboring HGs. **(C)** Chromosomal distribution of HGs genes across the CHM13 genome. Grey bars represent the total unique gene count per chromosome identified within ecDNA boundaries. Specific high-value cargo genes, including chromatin remodelers (e.g., *EZH2*, *KDM5A*) and transcription factors (e.g., *ZEB1*, *SOX9*), are highlighted and italicized to denote their functional potential within HGs.

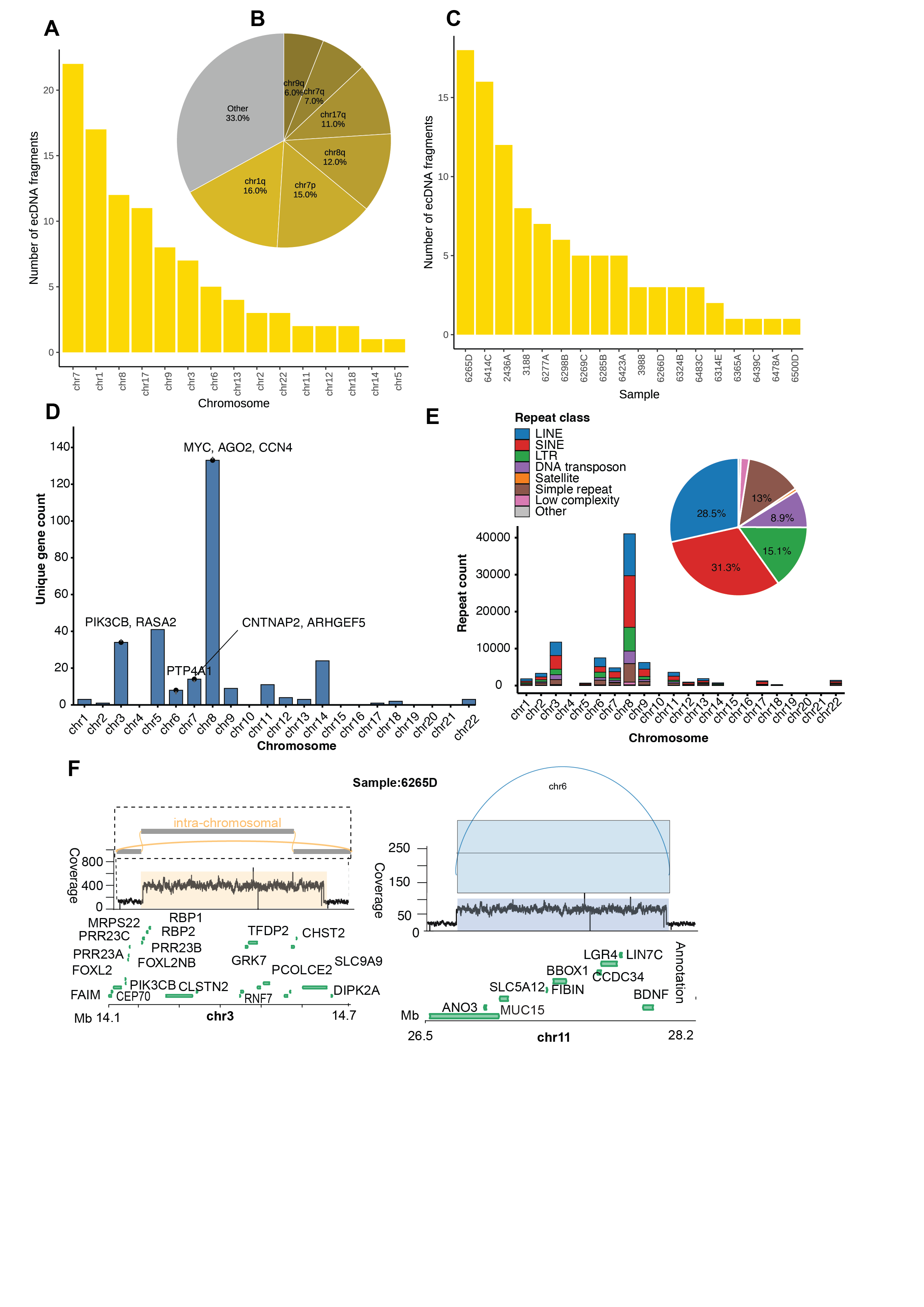

**Supplementary Figure 7. Chromosome-resolved ecDNA landscape and its relationship to telomere architecture in diffuse glioma.** **(A)** Bar plot summarizing ecDNA burden per chromosome across the cohort. ecDNA formation is highly uneven, with select chromosomes exhibiting disproportionate circular amplification activity, consistent with locus-specific structural fragility. **(B)** Arm-level circos plot depicting the proportional distribution of ecDNA across chromosome arms. Distinct arm-specific hotspots emerge, indicating focalized genomic regions that repeatedly undergo circularization and ecDNA generation across tumors. **(C)** Sample-level ecDNA burden plotted across patients. A subset of tumors shows markedly elevated ecDNA counts, capturing inter-tumoral variability in focal amplification severity. **(D)** Number of unique genes overlapping ecDNA intervals per chromosome, with labels indicating selected cancer- and ecDNA-relevant genes. **(E)** Chromosome-wise distribution of major repeat classes within ecDNA, shown as stacked bar plots of repeat annotation counts and whole-cohort proportional composition of major repeat classes across all ecDNA-overlapping repeat annotations. LINEs and SINEs were the most abundant repeat classes, and chromosome 8 exhibited the highest repeat contents. **(F)** In the representative ecDNA-positive sample **6265D**, a chimeric **intrachromosomal** assembly is shown, comprising multiple segments derived from the same chromosome and spanning regions that harbor oncogenic drivers (*left).* A simple circular ecDNA structure is also displayed, together with coverage tracks and gene annotations on the (*right).*

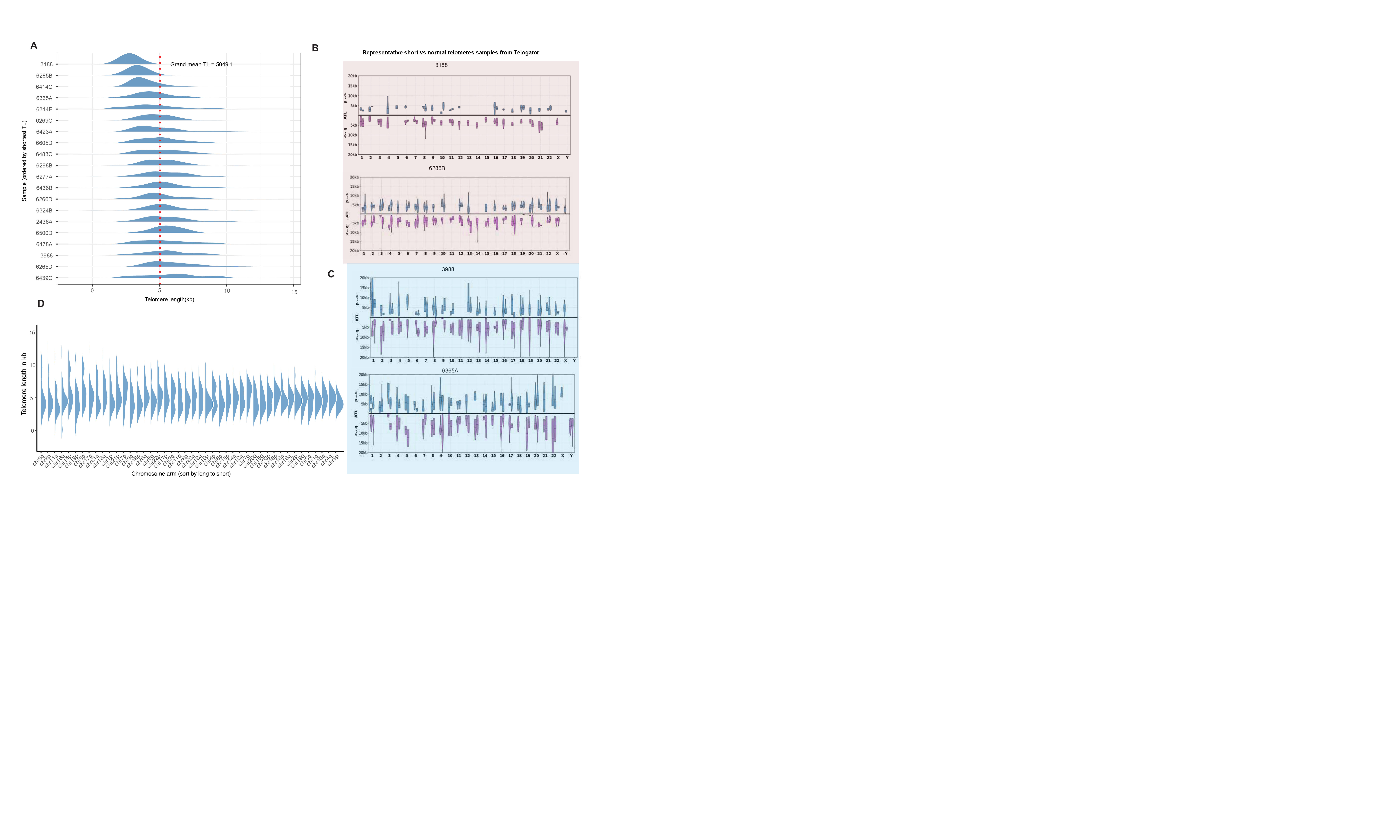

**Supplementary Figure 8. Telomere length landscapes across chromosome arms**. **(A)** Ridge plots illustrate the density distribution of absolute telomere lengths (kb) for individual samples (n =20). Each horizontal ridge corresponds to a single sample, with ridges ordered vertically by descending median length (longest at the bottom). The x-axis is oriented with the longest telomere lengths on the left, highlighting the progression toward telomere shortening on the right. The vertical red dashed line denotes the cohort-wide mean telomere length (5,049 bp). Peaks indicate the most frequent telomere lengths (modes) per sample, while extended tails represent subpopulations exhibiting significant telomere shortening or extreme lengthening. **(B–C)** Chromosome-arm telomere length profiles from Telogator for two representative tumors with short- and long-telomere states, shown separately for p- and q-arms within each chromosome. These examples illustrate pronounced arm-specific variation in telomere length across the astrocytoma genome. **(D)** Ridge plots illustrate the density distribution of absolute telomere lengths (kb) for individual chromosome arms across the study cohort. Arms are ordered vertically by descending median length, with the longest arms at the top and the shortest at the base. The peaks of each ridge indicate the most frequent telomere lengths (modes) within each arm, while the extended tails represent subpopulations exhibiting significant telomere shortening or extreme lengthening.

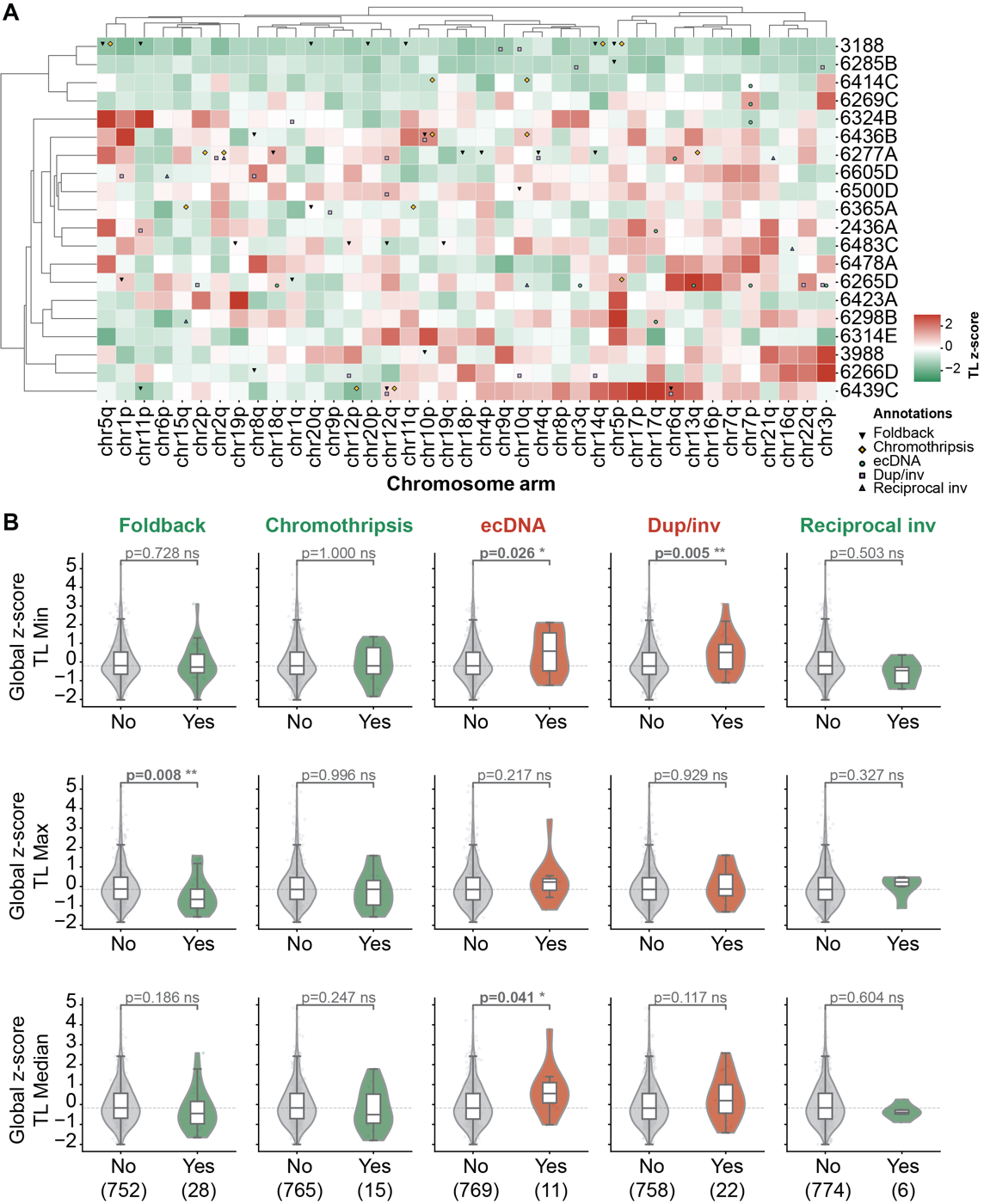

**Supplementary Figure 9. Telomere length heatmap and event-level associations. (A)** Clustered heatmap of globally z-scored median telomere length across 20 IDH-mutant astrocytoma samples (rows) and 39 non-acrocentric chromosome arms (columns). Color scale ranges from green (short TL, z < 0) through white to red (long TL, z > 0). Symbols overlaid on heatmap cells indicate the presence of specific genomic events: foldback inversions (triangle), chromothripsis (diamond), ecDNA (circle), duplicated inversions (square), and reciprocal inversions (triangle up). **(B)** Global z-scored telomere length distributions stratified by event presence for five primary signature events (columns) across three allelic TL measures (rows: TL Min, TL Max, TL Median). Violin plots compare arms without (grey) and with (colored) each event. P\-values from global permutation tests (10,000 permutations). Note that these univariate distributions include all event types regardless of statistical significance; the formal composite signature analysis (Figure 4D-F) provides the primary statistical model.

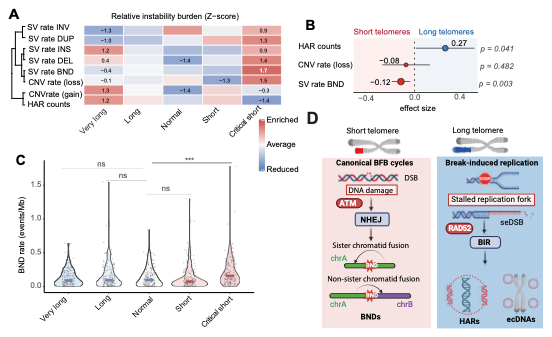

**Supplementary Figure 10. Schematic model of telomere shortening- and elongation-associated mechanisms of genome instability.** The model depicts two proposed pathways. In the setting of **telomere shortening**, telomere dysfunction induces DNA damage responses and promotes end-to-end fusion through **non-homologous end joining (NHEJ)** between sister or non-sister chromatids, triggering **breakage–fusion–bridge (BFB) cycles** and generating complex rearrangements enriched for **break-end (BND)** events. In contrast, under **telomere elongation/ALT-like states**, replication stress and stalled replication forks at these regions promote **break-induced replication** contribute to HGs and the emergence of **ecDNA**. Created with BioRender.

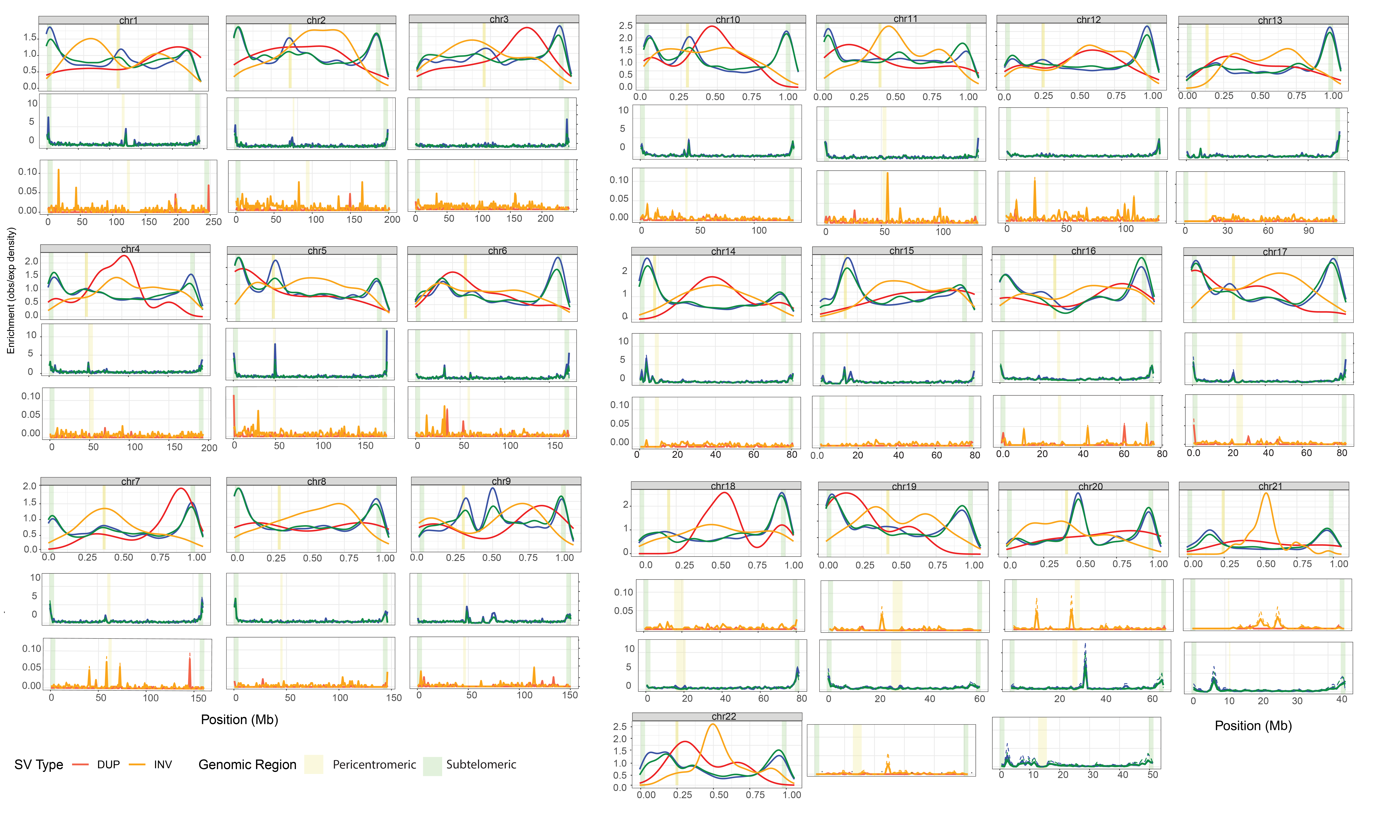

**Supplementary Figure 11. Chromosome-resolved landscape and regional enrichment of germline SV breakpoints.** Genome-wide germline SV breakpoints were mapped along each chromosome and scaled from 0 to 1 on the x-axis to represent the relative position from chromosome start to end. The y-axis denotes aggregated SV midpoint density across all individuals. Breakpoints are partitioned into centromeric regions (±5% around annotated centromeres; shaded khaki) and telomeric/subtelomeric regions (terminal 3% of each arm; shaded light green). This analysis reveals a pronounced accumulation of germline SV breakpoints near chromosome ends across the genome, with a secondary enrichment at centromeres. For each chromosome, normalized SV density per megabase is shown as line plots stratified by SV class (insertions, deletions, inversions, and duplications) across telomeric, centromeric, and interstitial regions. Deletions and insertions display the strongest telomeric enrichment, followed by moderate centromeric accumulation, whereas inversions show preferential localization to centromeric regions across most chromosomes. These patterns highlight conserved regional predispositions of the human genome to distinct germline SV mechanisms.

**Supplementary Figure 12. Chromosome-specific distribution and regional enrichment of somatic SV breakpoints (A)** Per chromosome, somatic SV breakpoints were aggregated across the cohort and projected onto a normalized chromosome coordinate system (x-axis scaled 0–1 from p-telomere to q-telomere), with the y-axis indicating SV-midpoint density. Telomeric and subtelomeric regions (terminal 3% plus adjacent 3% of each arm; light green) and centromeric zones (±5% around the centromere; khaki) are highlighted to delineate structurally vulnerable chromosomal boundaries. **(B)** Concordance between somatic and germline breakpoint enrichment by SV class. Scatter plots compare region-level SV densities for deletions and insertions (somatic, y-axis; germline, x-axis) across telomeric/subtelomeric, centromeric, and interstitial regions. Points are colored by genomic region, and fitted trend lines indicate a positive association between germline and somatic breakpoint landscapes, consistent with shared underlying fragility of telomeric and centromeric domains. Association strength and significance were assessed using Spearman correlation. (**C)** Trends represent the mean structural variant (BND) density (SVs/Mb) transitioning from centromeric to telomeric regions. Highlighted tracks identify specific chromosomes exhibiting a distinct "U-shaped" distribution, characterized by significantly higher breakpoint activity at the chromosomal extremities compared to interstitial regions, as assessed by a Wilcoxon rank-sum test.

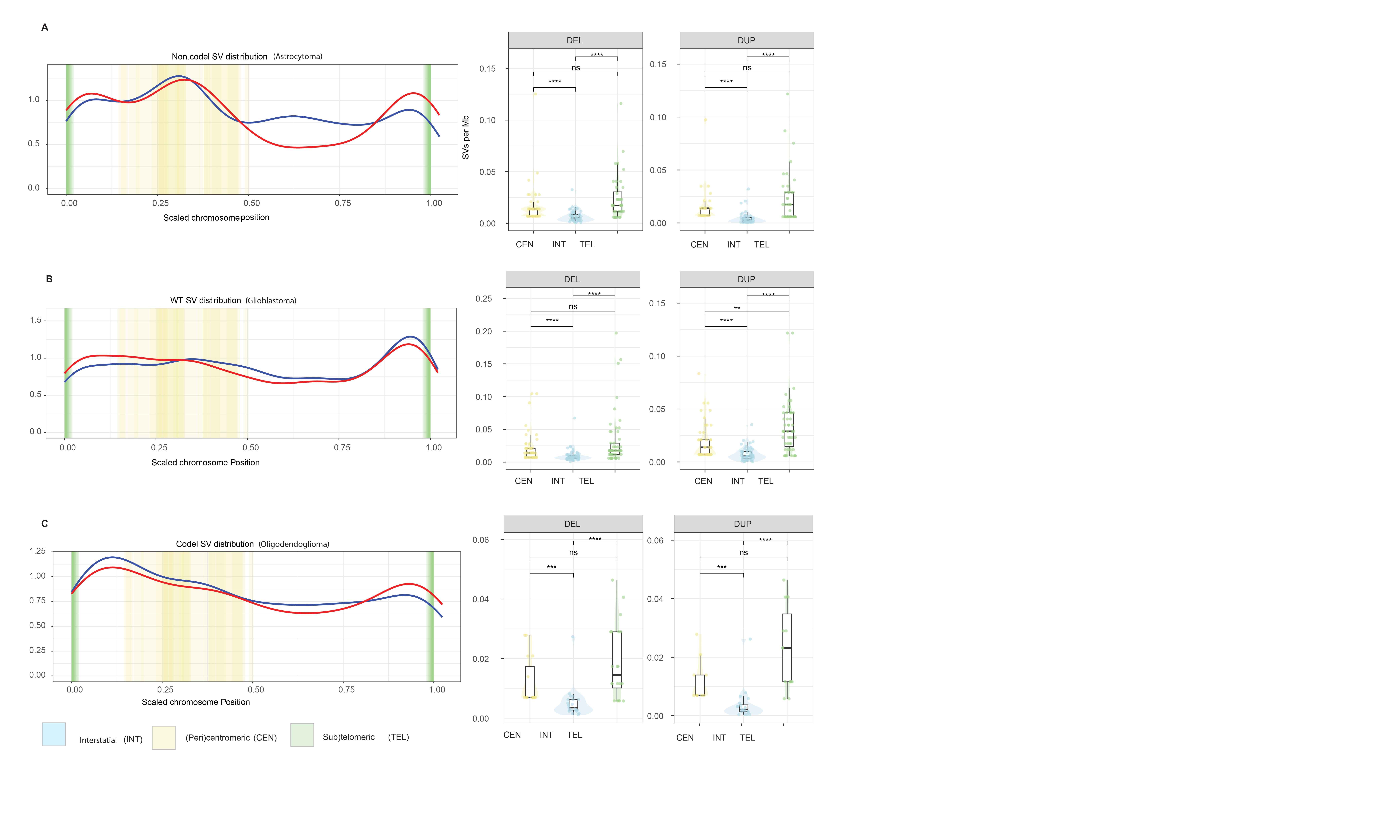

**Supplementary Figure 13. Chromosome-specific distribution and regional enrichment of structural variant breakpoints in public glioma cohorts (A–C)** Normalized SV densities (per megabase) were computed from publicly available glioma SV callsets (e.g., TCGA/GLASS-derived data) and mapped onto telomeric/subtelomeric, centromeric, and interstitial compartments for IDH-mutant astrocytomas, IDH-wild-type gliomas, and IDH-mutant oligodendrogliomas. SV densities were calculated after adjusting for chromosome-arm length and summarized within each subtype. Telomeric/subtelomeric regions were defined as the terminal 3% plus the adjacent 3% of each chromosome arm, and centromeric zones as ±5% around the centromere (see Methods). Statistical significance was assessed using a Wilcoxon rank-sum test, with significance denoted by asterisks where indicated.
